# TLNRD1 is a CCM complex component and regulates endothelial barrier integrity

**DOI:** 10.1101/2023.09.29.559344

**Authors:** Neil J. Ball, Sujan Ghimire, Gautier Follain, Ada O. Pajari, Monika Vaitkevičiūtė, Alana R. Cowell, Bence Berki, Johanna Ivaska, Ilkka Paatero, Benjamin T. Goult, Guillaume Jacquemet

## Abstract

We previously identified talin rod domain-containing protein 1 (TLNRD1) as a potent actin-bundling protein in vitro. Here, we report that TLNRD1 is primarily expressed in the vasculature in vivo and that its depletion leads to vascular abnormalities in vivo and loss of barrier integrity in cultured endothelial cells. We demonstrate that TLNRD1 is a component of the cerebral cavernous malformations (CCM) complex through its direct, high-affinity interaction with CCM2. Modeling and functional testing of TLNRD1 and CCM2 mutants reveal that their interaction is mediated by a hydrophobic C-terminal helix in CCM2 that attaches to a hydrophobic groove on the 4-helix domain of TLNRD1. Disruption of this binding interface leads to CCM2 and TLNRD1 accumulation in the nucleus and actin fibers. Notably, a CCM2 pathogenic mutation linked to vascular dementia in patients maps to the interface and disrupts the interaction. Our findings indicate that CCM2 controls TLNRD1 localization to the cytoplasm and inhibits its actin-bundling activity. Based on these results, we propose a new pathway by which the CCM complex modulates the actin cytoskeleton and vascular integrity by controlling TLNRD1 bundling activity.

## Introduction

The actin cytoskeleton is essential for most cellular processes, including cell movement, cell division, and cell shape maintenance. Consequently, aberrant actin cytoskeleton regulation is linked to multiple illnesses, including cancer and immunological and neurological disorders. Actin dynamics are regulated by various proteins, which include nucleators, polymerizers, depolymerizers, and crosslinkers, all working together to ensure spatially and temporally appropriate assembly of actin structures (Lappalainen et al., 2022). While the role of individual actin regulatory proteins is starting to be understood in cells, their functions in living organisms often remain elusive.

We previously identified talin rod domain-containing protein 1 (TLNRD1, also known as MESDC1) as a potent actin-bundling protein in vitro and cultured cells (Cowell et al., 2021). We found that TLNRD1 is homologous to the R7R8 domains of the cytoskeletal adaptor talin (Gingras et al., 2010) and exists as a constitutive homodimer (Cowell et al., 2021). By solving the TLNRD1 structure, we demonstrated that TLNRD1 comprises a 4-helix domain (TLNRD1^4H^, equivalent to talin R8) inserted into a 5-helix domain (TLNRD1^5H^, equivalent to talin R7) to form a 9-helix module and that TLNRD1 homodimerization is mediated via an interface located on the 4-helix module. Like talin R7R8, TLNRD1 binds F-actin, but because TLNRD1 forms an antiparallel dimer, it also bundles F-actin. In cancer cells, TLNRD1 localizes to the cytoplasm and accumulates on actin bundles and in filopodia (Cowell et al., 2021). Functionally, we reported that TLNRD1 expression enhanced filopodia formation and cancer cell migration, while TLNRD1 down-regulation had the opposite effect. Other studies have shown that TLNRD1 overexpression is associated with increased proliferation and xenograft growth in hepatocellular carcinoma (Tatarano et al., 2012; Wu et al., 2017) and that TLNRD1 depletion reduced bladder cancer cell migration and invasion (Nagy et al., 2018). While TLNRD1 appears to modulate cancer cell functions, little is known about the physiological functions of TLNRD1 in normal tissue.

The cerebral cavernous malformations (CCM) complex is a trimeric protein assembly critical for vascular homeostasis. This complex comprises three proteins: Krev interaction trapped protein 1 (KRIT1 or CCM1), Cerebral cavernous malformations 2 protein (CCM2), and Programmed cell death protein 10 (PDCD10 or CCM3). Together, they orchestrate endothelial cell functions by modulating endothelial cell junctions and the actin cytoskeleton. The CCM complex is pivotal in regulating the MEKK3-MEK5-ERK5 signaling cascade and the small GTPase RhoA (Su and Calderwood, 2020). Specifically, by regulating RhoA activity, the CCM complex facilitates actin remodeling, indispensable for effective cell migration and underpins endothelial cell junction robustness. Importantly, disruptions in the functions of these proteins, whether through mutations or loss of function, are intrinsically linked to CCM disease, a neurovascular disorder characterized by the emergence of blood-filled cavernomas within the central nervous system (Riolo et al., 2021). Here, we report that TLNRD1 is a member of the CCM complex. In zebrafish embryos, silencing of TLNRD1 results in vascular malformations, while in endothelial cells, it disrupts monolayer integrity and actin organization. We found that TLNRD1 directly binds to CCM2 through a hydrophobic interaction involving a C-terminal helix in CCM2 and a groove on TLNRD1’s 4-helix domain. As our findings demonstrate that CCM2 inhibits TLNRD1’s actin-binding activity, we propose that the CCM complex can also modulate the actin cytoskeleton and vascular integrity by controlling TLNRD1 activity.

## Results

### TLNRD1 is a putative member of the CCM complex

We previously described TLNRD1 as an actin-bundling protein contributing to filopodia formation in cancer cells (Cowell et al., 2021). However, the broad localization of TLNRD1 in cells suggests that it likely has other roles as well. To further understand TLNRD1 cellular functions, we sought to identify TLNRD1 binding partners using an unbiased massspectrometry approach. We performed GFP pulldowns from cells expressing either GFP or GFP-TLNRD1 followed by mass spectrometry analysis (Fig. 1A and Fig. 1B). Using this strategy, we identified 89 proteins as specifically enriched to TLNRD1 pulldowns (Fig. 1A and 1B and Table S1). Interestingly, mapping putative TLNRD1 binders onto a protein-protein interaction network revealed that several hits, namely KRIT1 (also known as CCM1), CCM2, PDCD10 (also known as CCM3), serine/threonine kinase 25 (STK25), and integrin *β*1-binding protein 1 (ITGB1BP1, also known as ICAP1) cluster together (Fig. 1C). These proteins are known to assemble the CCM complex, a trimeric protein complex of KRIT1-CCM2-PDCD10, with STK25 and ITGB1BP1 being accessory members (Yang et al., 2023; Faurobert et al., 2013). In addition, western blot analyses confirmed that KRIT1 and ITGB1BP1 co-purify with GFP-TLNRD1, further validating our mass-spectrometry analyses (Fig. 1D). These results led us to speculate that TLNRD1 could be a full, or accessory, member of the CCM complex.

**Fig. 1.**
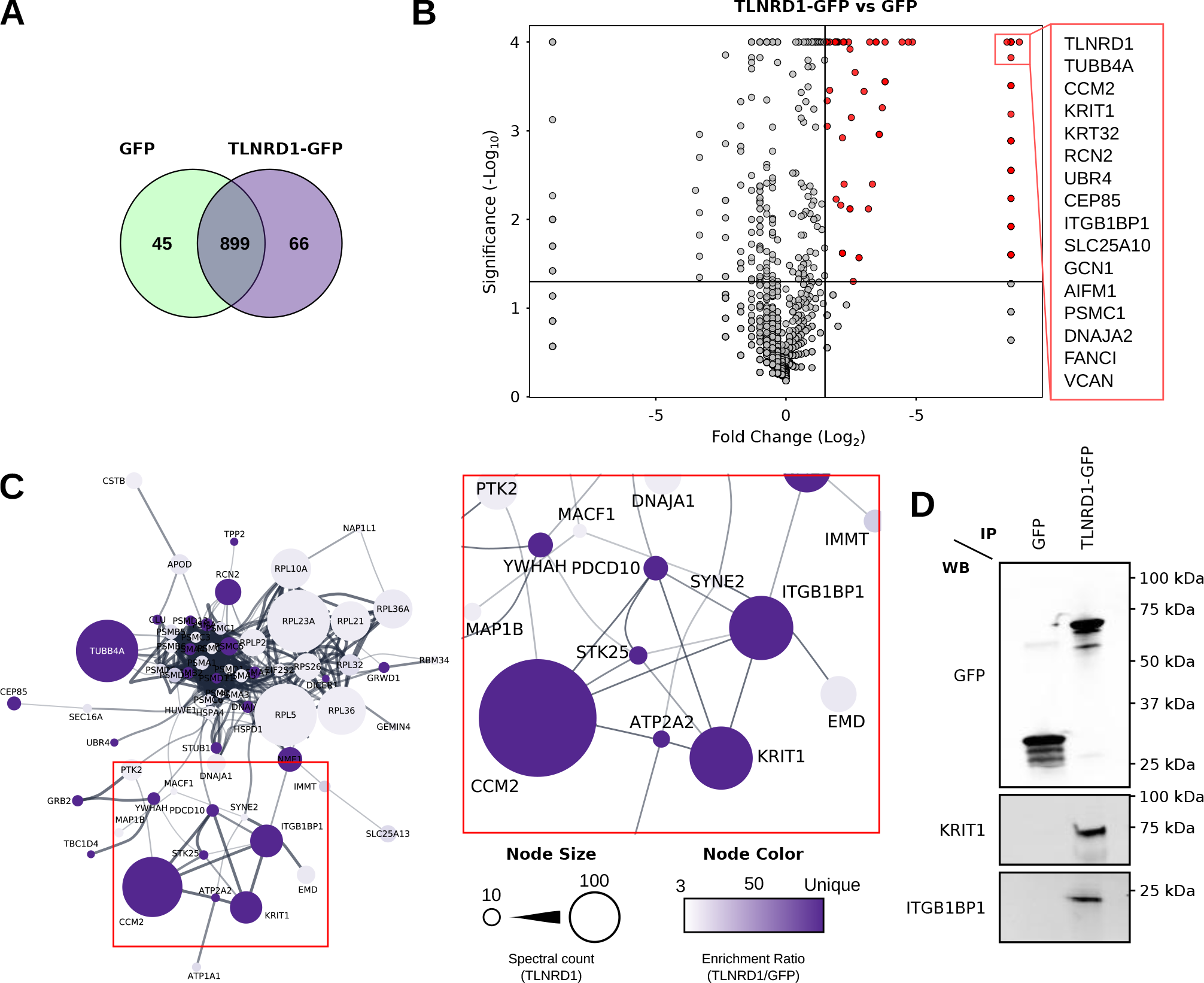
Mass spectrometry analyses identify TLNRD1 as a putative member of the CCM complex. (**A**-**C**) Mass spectrometry (MS) analysis of GFP-TLNRD1-binding proteins. Comparison of the GFP-TLNRD1 and GFP datasets are displayed as a Venn diagram (**A**) and as a volcano plot (**B**). In the volcano plot, the enrichment ratio (TLNRD1 over GFP) for each protein detected is plotted against the significance of the association (see Table S1 for the MS data). (**C**) Proteins specifically enriched to TLNRD1 were mapped onto a protein-protein interaction network (STRING, see the Materials and Methods for details). Each node (circle) represents a protein (labeled with gene name), and each edge (line) represents a reported interaction between two proteins. The node’s color indicates the enrichment ratio of that particular protein (TLNRD1 over GFP). The node’s area represents the spectral count of that specific protein in the TLNRD1-GFP dataset. (**D**) GFP-pulldown in HEK293T cells expressing GFP-TLNRD1 or GFP alone. KRIT1 and ITGB1BP1 recruitment to the bait proteins was then assessed by western blotting (representative of three biological repeats).

We previously showed that TLNRD1 and ITGB1BP1 accumulate at the tips of myosin-X (MYO10)-induced filopodia in cancer cells (Jacquemet et al., 2019; Cowell et al., 2021). Therefore, we next assessed, using structured illumination microscopy, the ability of the other CCM complex members to localize to filopodia tips (Fig. S1). These experiments revealed that CCM2 and PDCD10 could also be found at the tip of MYO10-induced filopodia (Fig. S1), further linking TLNRD1 to the CCM complex.

### TLNRD1 depletion leads to vascular malformation in vivo

While the individual CCM complex components are expressed in diverse cell types, the CCM complex has a well-documented role in forming and maintaining the vasculature. Indeed, loss of function mutations of KRIT1, CCM2, or PDCD10 leads to cerebral cavernous malformations, which are vascular lesions defined in patients by blood-filled endothelial cell caverns and an absence of a mature vessel wall (Chohan et al., 2019).

Analysis of publicly available single-cell RNA-Seq datasets (Tabula Muris, (Schaum et al., 2018)) revealed that in adult mice, TLNRD1 is expressed in endothelial cells in the brain (Fig. 2A). TLNRD1 expression is, however, not limited to the brain vasculature and TLNRD1 transcripts were also detected in other organs, for instance, in the endothelial cells in the heart as well as other cell types including fibroblasts and leukocytes (Fig. 2B). To validate these datasets, we stained and imaged mouse brain slices and observed that TLNRD1 was expressed in PECAM-positive vessels (Fig. 2C). Together these data indicate that TLNRD1 is expressed in the vascular endothelium in vivo.

**Fig. 2.**
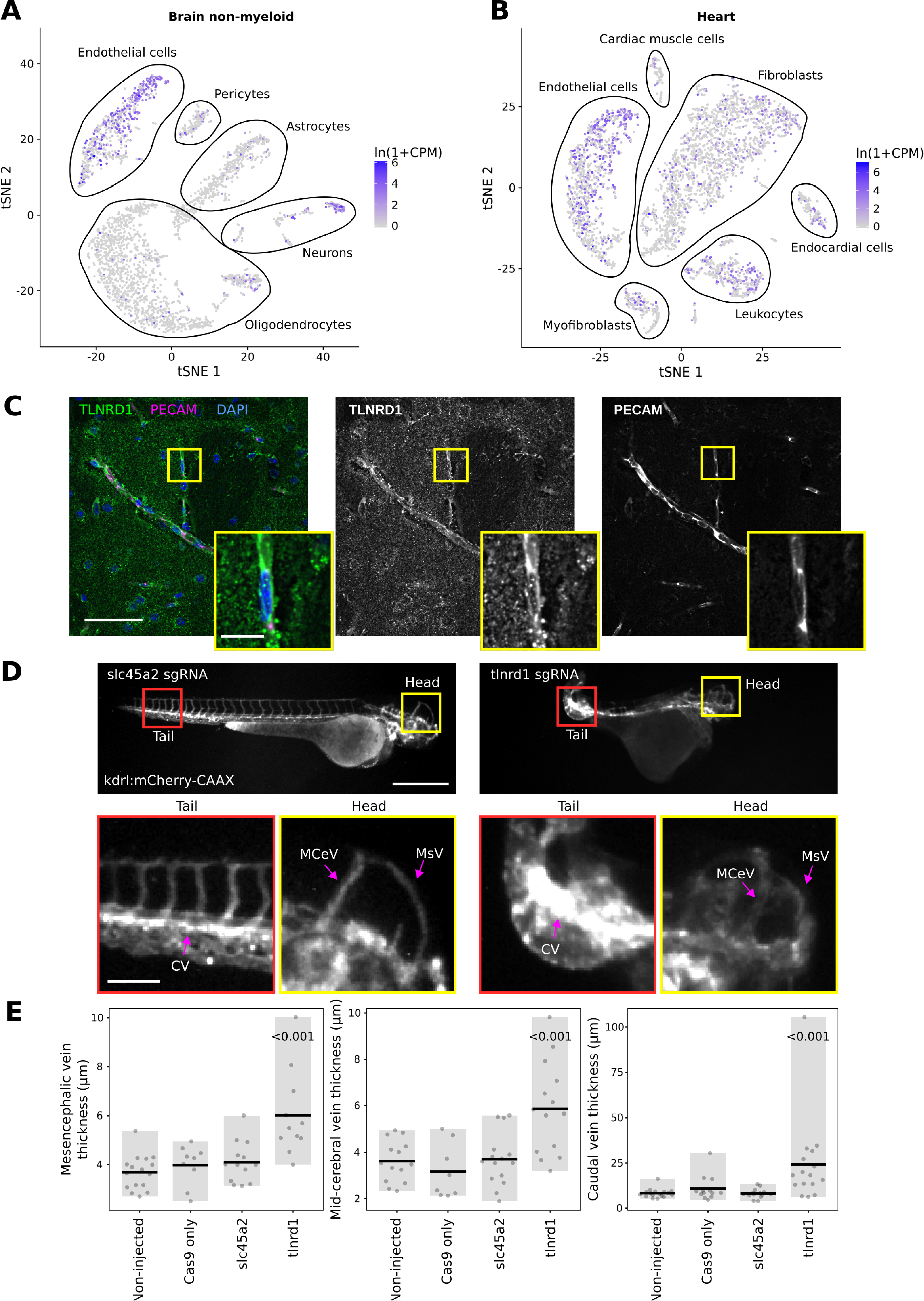
TLNRD1 is expressed in endothelial cells in vivo and regulates the vascular system. (**A, B**): TLNRD1 expression in mouse brain (**A**) and mouse heart (**B**). This single-cell RNA sequencing data is from the Tabula Muris dataset (Schaum et al., 2018). (**C**): Mouse brain slices were stained for TLNRD1, PECAM, and DAPI and imaged using a spinning disk confocal microscope. A single Z-plane is displayed. The yellow square highlights a magnified region of interest (ROI). Scale bars: (main) 50 µm and (inset) 10 µm. (**D, E**): kdrl:mCherry-CAAX zebrafish embryos were injected with recombinant Cas9 alone or together with sgRNA targeting TLNRD1 or slc45a2. The embryos were then imaged using a fluorescence microscope. (**D**) Representative images are displayed. The yellow and red squares highlight ROIs, which are magnified. The mesencephalic (MsV), mid-cerebral (MCeV), and caudal (CV) vein plexus are highlighted. Scale bars: (main) 500 µm and (inset) 100 µm. (**E**) The thickness of the mesencephalic, mid-cerebral, and caudal vein plexus measured from microscopy images are plotted as dot plots (non-injected, n=17; Cas9, n=13; slc45a2, n=14; TLNRD1, n=16). The grey bar highlights the data distribution, while the black line indicates the mean. The p-values were determined using a randomization test.

Zebrafish embryos are robust model organisms for studying the cardiovascular system (Hogan and Schulte-Merker, 2017). In addition, mutation of CCM complex components, including KRIT1, CCM2, or PDCD10, in zebrafish embryos leads to defects in the vasculature (Yoruk et al., 2012; Hogan et al., 2008). Therefore, we next investigated the impact of TLNRD1 gene disruption in zebrafish embryos using CRISPR (Fig. S2A). Strikingly, zebrafish embryos treated with anti-TLNRD1 CRISPR guide RNAs exhibited severe abnormal vascular morphology at several anatomical sites. In particular, the mesencephalic, mid-cerebral, and caudal vein plexus were dilated in TLNRD1-targeted embryos (Fig. 2D and 2E). These phenotypes were not observed in the control embryos. TLNRD1-targeted embryos also had lower heart rates than control embryos, but these results did not reach statistical significance (Fig. S2B). Interestingly, single-cell RNA-Seq datasets revealed that TLNRD1 expression peaks at 24 h post-fertilization during zebrafish embryo development, where it is principally expressed in the blood vasculature (Zebrahub (Lange et al., 2023)). Similarly, TLNRD1 is also expressed in the vasculature in human embryos (Fig. S2C, (Xu et al., 2023)). Importantly, reanalysis of publicly available datasets revealed that TLNRD1 expression is up-regulated in CCM lesions from patients (Fig. S2D, Subhash et al., 2019). Altogether, our data point toward a crucial, previously unknown role for TLNRD1 in regulating the vasculature in vivo.

### TLNRD1 modulates junctional integrity in endothelial cells

Zebrafish embryos are powerful tools to study and observe the development of the cardiovascular system; however, they are less amenable to mechanistic studies. Therefore, we next investigated the contribution of TLNRD1 to endothelial cell functions. Human umbilical vein endothelial cells (HUVEC) express TLNRD1 (Fig. S3A) (Cowell et al., 2021). In HUVEC monolayers, TLNRD1 localized to the cytoplasm and accumulated on actin bundles, including stress fibers and filopodia (Fig. 3A). TLNRD1 also localized on the actin structures near cell-cell junctions (Fig. 3A).

**Fig. 3.**
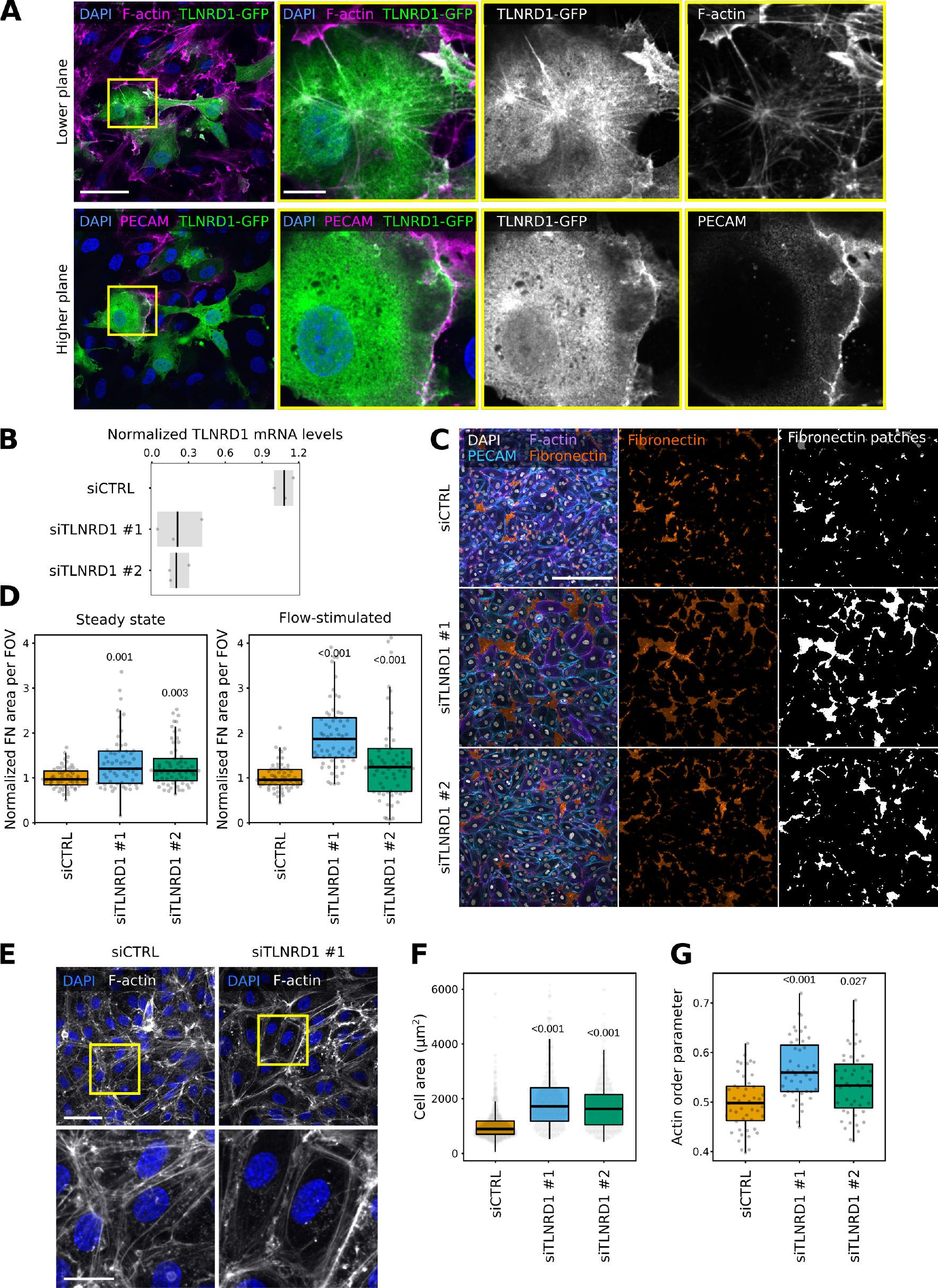
TLNRD1 loss disrupts endothelial monolayer integrity. (**A**) HUVECs expressing TLNRD1-GFP were fixed, stained for DAPI, F-actin, and PECAM, and imaged using a spinning disk confocal microscope. Two Z-planes from the same field of view are displayed. The yellow squares highlight magnified ROIs. Scale bars: (main) 50 µm and (inset) 10 µm. (**B**-**D**) TLNRD1 expression was silenced in HUVECs using two independent siRNA. (**B**) TLNRD1 expression levels were determined by qPCR. (**C**-**D**) HUVEC cells were allowed to form a monolayer in the presence or absence of flow stimulation. Cells were then fixed and stained for DAPI, F-actin, PECAM, and Fibronectin (without permeabilization) before imaging on a spinning disk confocal microscope. (**C**) Representative maximum intensity projections are displayed (flow stimulation). Scale bar: 250 µm. (**D**) The area covered by fibronectin patches in each field of view was then quantified (3 biological repeats, n >60 fields of view per condition). (**E**-**G**) TLNRD1 expression was silenced in HUVECs using two independent siRNAs, and cells were allowed to form a monolayer without flow stimulation. Cells were then fixed and stained for DAPI and F-actin. Images were acquired using a spinning disk confocal microscope. (**E**) Representative SUM projections are displayed. Scale bar: (main) 50 µm and (inset) 20 µm. (**F**) The cell area was measured using manual cell segmentation (3 biological repeats, >45 fields of view, >460 cells per condition). (**G**) The actin organization (order parameter) was quantified using Alignment by Fourier Transform (Marcotti et al., 2021). The results are shown as Tukey boxplots. The whiskers (shown here as vertical lines) extend to data points no further from the box than 1.5× the interquartile range. The P-values were determined using a randomization test.

Next, we assessed the contribution of TLNRD1 to endothelial monolayer integrity. TLNRD1 expression was silenced using two independent siRNAs (Fig. 3B). After three days, the resulting monolayers were fixed, stained for fibronectin without permeabilization, and imaged. This approach allowed us to assess the permeability of the endothelial monolayer by quantifying the size and number of fibronectin patches, as these patches on the ventral side of the monolayer become accessible to the antibody when the junction above them is leaky (Fig. S3B). Importantly, these experiments revealed that TLNRD1 silencing disrupted the monolayer, both in the presence or absence of flow stimulation (Fig. 3C, 3D, and Fig. S3C). When imaged at higher magnification, we observed that TLNRD1-silenced cells were more spread out than control cells (Fig. 3E and 3F). In addition, the overall organization of the actin cytoskeleton in the formed monolayers appeared altered. In TLNRD1-silenced cells, actin stress fibers were more prominent on individual cells’ edges, rendering the overall actin organization in the monolayer more organized (Fig. 3E and 3G). Altogether, our data indicate that TLNRD1 contributes to overall actin cytoskeleton organization and junctional integrity in endothelia.

### TLNRD1 binds to CCM2 via its 4-helix bundle

Having shown that TLNRD1 co-purifies with the CCM complex (Fig. 1) and that TLNRD1 depletion leads to vascular phenotypes in both zebrafish embryos (Fig. 2) and endothelial cells (Fig. 3), we next wanted to map the interaction(s) between TLNRD1 and the CCM complex. We implemented a protein trapping strategy to identify which CCM protein(s) interacts with TLNRD1. CCM2 and PDCD10 were targeted to the mitochondria, and the ability of TLNRD1 to be recruited to this compartment was then analyzed using fluorescence microscopy (Fig. 4A and 4B). Using this strategy, we found that TLNRD1 strongly colocalized with mitochondrial-targeted (mito)-CCM2 but not mito-PDCD10 (Fig. 4A and 4B). Interestingly, TLNRD1 and mito-CCM2 clustered strongly together, forming aggregates (Fig. 4A, (Cowell et al., 2021)). Significantly, the recruitment of TLNRD1 to mito-CCM2 was not affected when KRIT1 expression was silenced using siRNA (Fig. S4B and S4C), indicating that the TLNRD1-CCM2 interaction does not require KRIT1. Next, we expressed TLNRD1 and CCM2 in endothelial cells. We found that both proteins colocalize on actin structures and in the cytosol (Fig. 4C). Our data indicate that TLNRD1 interacts with the CCM complex via CCM2.

**Fig. 4.**
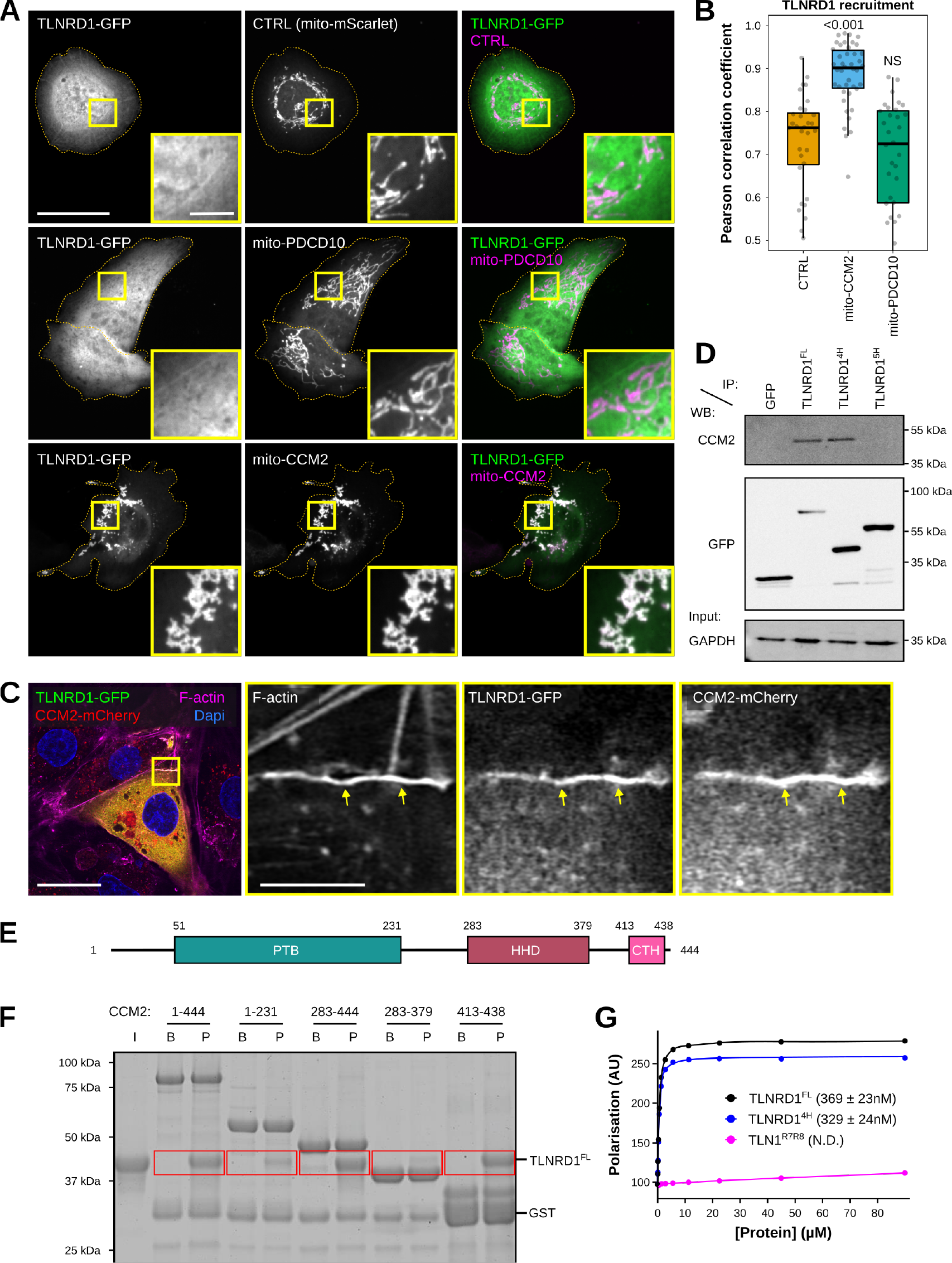
The TLNRD1-CCM2 interaction involves the TLNRD1 4-helix bundle and a C-terminal helix in CCM2. (**A**-**B**) U2OS cells expressing TLNRD1-GFP and mitomScarlet (CTRL), mito-PDCD10-mScarlet, or mito-CCM2-mScarlet were imaged using a spinning disk confocal microscope. (**A**) Representative single Z-planes are displayed. Dashed yellow lines highlight the cell outlines. The yellow squares highlight magnified ROIs. Scale bars: (main) 25 µm and (inset) 5 µm. (**B**) 3D colocalization analysis was performed using the JACoP Fiji plugin (three biological repeats, n> 31 image stacks per condition). The results are shown as Tukey boxplots. The whiskers (shown here as vertical lines) extend to data points no further from the box than 1.5× the interquartile range. The P-values were determined using a randomization test. NS indicates no statistical difference between the mean values of the highlighted condition and the control. (**C**) HUVECs expressing TLNRD1-GFP and CCM2-mCherry were stained for F-actin and Dapi and imaged using an Airyscan confocal microscope. A single Z-plane is displayed. The yellow squares highlight a magnified ROI. Scale bars: (main) 25 µm and (inset) 5 µm. (**D**) GFP-pulldown in HEK293T cells expressing GFP-TLNRD1, GFP-TLNRD1^4H^. GFP-TLNRD1^5H^ or GFP alone. CCM2 recruitment to the bait proteins was assessed by western blotting (representative of three biological repeats). (**E**): CCM2 schematic showing the boundaries of the phosphotyrosine binding (PTB) domain, the harmonin homology domain (HHD), and the C-terminal helix (CTH). (**F**) A GST-pulldown assay was used where Glutathione agarose-bound GST-CCM2 fragments (beads: B) were incubated with recombinant TLNRD (input: I). After multiple washes, proteins bound to the beads (pellet: P) were eluted. A representative gel of three independent repeats is displayed. Red boxes highlight areas of interest in the gel. (**G**) A fluorescence polarization assay was used to determine the Kd of the interaction between TLNRD1, TLNRD1^4H^, or TLN1^R7R8^ with SUMO-CCM2^CTH^. Kd values (nM) are shown in parentheses. ND, not determined.

To map the TLNRD1 domain interacting with CCM2, we performed GFP-trap experiments in cells expressing GFP, GFP-TLNRD1, GFP-TLNRD1^4H^, or GFP-TLNRD1^5H^ (Fig. 4D). CCM2 co-precipitated with TLNRD1 and TLNRD1^4H^ but not TLNRD1^5H^, indicating that CCM2 interacts with TLNRD1 via the TLNRD1 4-helix bundle (Fig. 4D). Next, we used a GST-pulldown assay with recombinant proteins to determine whether the TLNRD1-CCM2 interaction is direct (Fig. S4D). GST-CCM2 co-purified with recombinant TLNRD1 and TLNRD1^4H^ but not with TLN1^R7R8^ (Fig. S4D), despite TLN1^R7R8^ and TLNRD1^4H^ sharing a similar domain organization and structure (Cowell et al., 2021). Our data indicate that TLNRD1 directly interacts with CCM2 via the TLNRD1 4-helix bundle domain.

### The TLNRD1-CCM2 interaction involves the C-terminal helix in CCM2

CCM2 contains two characterized domains, an N-terminal phosphotyrosine binding (PTB) domain and a C-terminal harmonin homology domain (HHD) (Fig. 4E). To map the region of CCM2 responsible for interacting with TLNRD1, we generated several CCM2 truncated constructs designed based on the known CCM2 structures (Fisher et al., 2015; Wang et al., 2015; Fisher et al., 2013) and the AlphaFold CCM2 structure prediction (Jumper et al., 2021). Using GST-pulldown and recombinant proteins, we found that TLNRD1 co-purifies in vitro with CCM2, CCM2^283-444^, and CCM2^413-438^ but not with CCM2^1-231^ or CCM2^283-379^ (Fig. 4F). A Fluorescence Polarization (FP) assay revealed that TLNRD1 and TLNRD1^4H^ interact with CCM2^413-438^ with a dissociation constant (Kd) in the nanomolar range (Fig. 4G). At the same time, no interaction between CCM2^413-438^ and TLN1^R7R8^ was detected (Fig. 4G). Our data indicate that CCM2 does not bind TLNRD1 via the previously characterized CCM2 HHD or PTB domains. Instead, CCM2 interacts with TLNRD1 via the CCM2^413-438^ region, which AlphaFold predicts as a helix, which we call here the CCM2 C-terminal helix (CTH) for simplicity.

### The TLNRD1-CCM2 binding interface involves a hydrophobic groove on TLNRD1 and hydrophobic residues of CCM2

Having mapped the regions in CCM2 and TLNRD1 responsible for their interaction, we next modeled the CCM2-TLNRD1 complex using ColabFold (Mirdita et al., 2022) (Fig. 5A). The ColabFold prediction suggests that CCM2^CTH^ binds to TLNRD1^4H^, with each TLNRD1 monomer capable of binding one CCM2^CTH^ (Fig. 5A and Movie 1) in agreement with our biochemical analysis (Fig. 4H). Analysis of the predicted binding interface indicated that the TLNRD1-CCM2 interaction was predominantly hydrophobic, with the hydrophobic residues of the amphipathic CCM2^CTH^ inserted into a hydrophobic groove on the surface of TLNRD1^4H^ (Fig. 5B). To test this model, we designed targeted point mutations in TLNRD1 and in CCM2 to perturb their interaction. Two CCM2 mutants were designed, namely CCM2^I432D^ and CCM2^W421D/D422A^ (termed here CCM2^WD/AA^). Using the FP assay, we found that the CCM2^WD/AA^ and CCM2^I432D^ mutations abolish the TLNRD1-CCM2^CTH^ interaction in vitro validating our structural modeling and the critical importance of the hydrophobic interface in the interaction (Fig. 5D and 5E). Interestingly, an I428S mutation in CCM2 has been reported in a patient diagnosed with vascular dementia (CCM2^I428S^; (Mönkäre et al., 2021)) and this isoleucine is on the same face of the CTH helix as the mutations we designed, so we predicted it too would impact the TLNRD1-CCM2 binding interface. Testing the CCM2^I428S^ mutant in the FP assay showed that it had a similar effect to the CCM2^WD/AA^ and CCM2^I432D^ mutations, disrupting the TLNRD1-CCM2^CTH^ interaction (Fig. 5D).

**Fig. 5.**
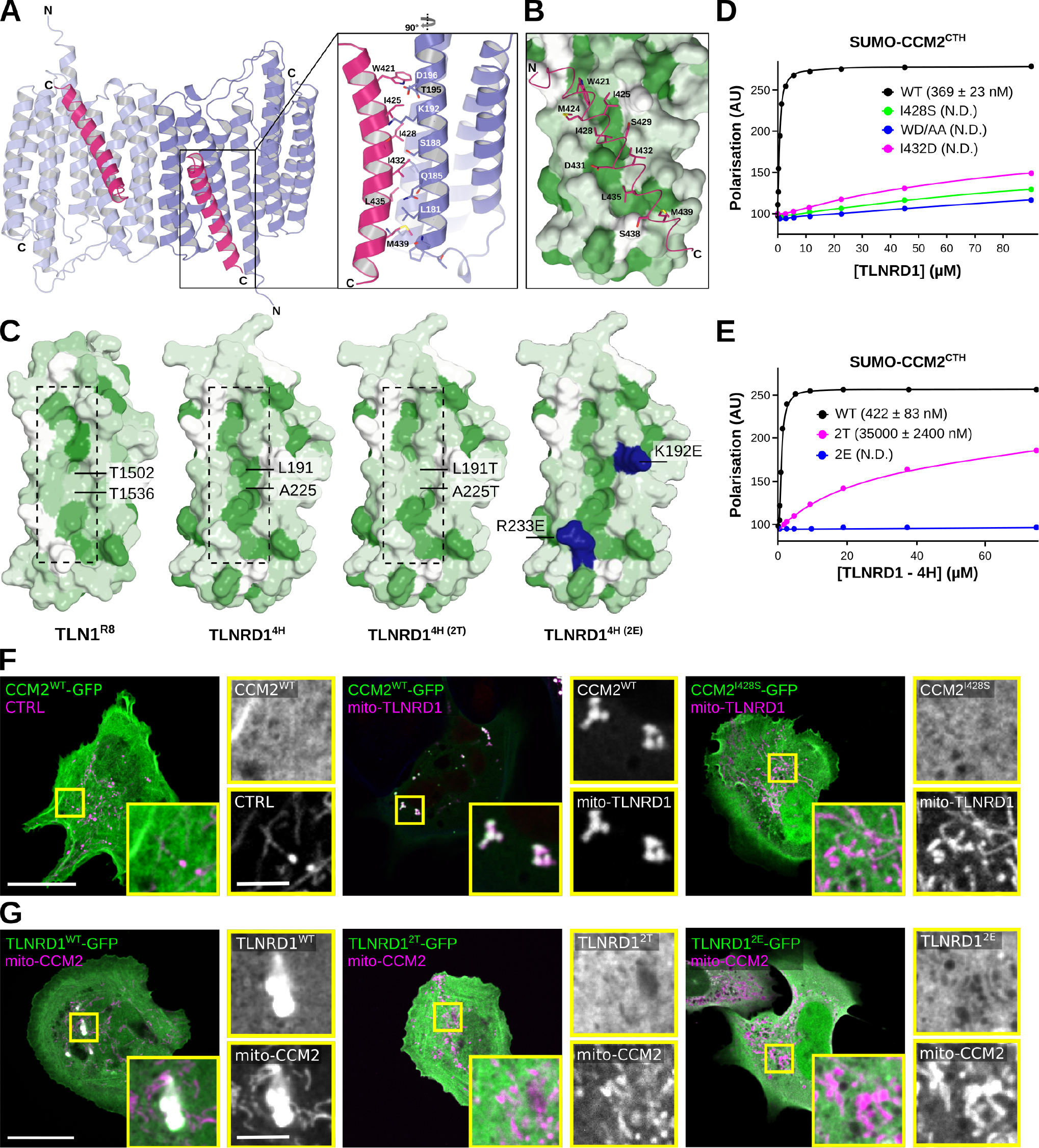
The TLNRD1-CCM2 binding interface involves a hydrophobic groove on TLNRD1 and hydrophobic residues of CCM2. (**A**-**B**) Modeling of the TLNRD1-CCM2 complex using ColabFold using the TLNRD1 crystal structure (PDB accession code 6XZ4) as a template. (**A**) Overall view of the predicted complex. The TLNRD1 monomers are colored blue, and the CCM2^CTH^ helices are colored pink. The TLNRD1-CCM2 binding area is magnified, and the residues contributing to the interface are shown as sticks. (**B**) TLNRD1 has a hydrophobic channel (green) on the surface, which could facilitate CCM2 (pink) binding. TLNRD1^4H^ was colored by hydrophobicity using the AA index database (entry FASG890101, (Nakai et al., 1988)) in PyMOL, where green denotes hydrophobic residues and white polar residues. CCM2^CTH^ is shown as sticks and predominantly contacts the hydrophobic region on TLNRD1^4H^. (**C**) Comparison of the hydrophobic channel on the surface of TLNRD1^4H^ and the equivalent region on TLN1^R8^. The TLNRD1^2T^ mutant was designed to mimic the surface of TLN1^R8^. The green color denotes hydrophobic residues. On the TLNRD1^2E^, the mutated basic residues are highlighted in blue. (**D**) Fluorescence polarization was used to determine the Kd of the interaction between TLNRD1 and various SUMO-CCM2^CTH^ constructs (WT, I428S, I432D, and W412A/D422A). Kd values (nM) are shown in parentheses. ND, not determined. (**E**) Fluorescence polarization was used to determine the Kd of the interaction between CCM2^CTH^ and various TLNRD1^4H^ constructs (WT, 2T, and 2E). Kd values (nM) are shown in parentheses. ND, not determined. (**F**) U2OS cells expressing various GFP-tagged CCM2 constructs and mito-TLNRD1-mScarlet or mito-mScarlet (CTRL) were imaged using a spinning disk confocal microscope. Representative single Z-planes are displayed. See also Fig S4E and 4F. Scale bars: (main) 25 µm and (inset) 5 µm. (**G**) U2OS cells expressing various GFP-tagged TLNRD1 constructs and mito-CCM2-mScarlet or mito-mScarlet (CTRL) were imaged using a spinning disk confocal microscope. Representative maximum intensity projections are displayed. Scale bars: (main) 25 µm and (inset) 5 µm.

We next designed mutations in TLNRD1 aiming at disrupting CCM2 binding; these double mutations, TLNRD1^L191T/A225T^ (termed here TLNRD1^2T^) and TLNRD1^K192E/R233E^ (termed here TLNRD1^2E^) were introduced into TLNRD1^4H^. Importantly, the TLNRD1^2T^ mutant was designed to mimic the surface of TLN1^R8^ (Fig. 5C), which does not bind to CCM2 (Fig. 4G). We found that the TLNRD1^2E^ abolished binding to CCM2^CTH^ (Fig. 5E), and the TLNRD1^2T^ mutant reduced the TLNRD1 affinity for CCM2 by 150-fold (Fig. 5E). Next, we validated our in vitro experiments in cells using the previously described mitochondrial trapping strategy. TLNRD1 or CCM2 was targeted to the mitochondria, and the ability of various TLNRD1 and CCM2 constructs to be recruited to this compartment was then analyzed using fluorescence microscopy (Fig. 5G). As expected, we found that CCM2 strongly colocalized with mito-TLNRD1 and that TLNRD1 strongly colocalized with mito-CCM2 (Fig. 5F and 5G). However, deletion of CCM2^CTH^ (CCM2^ΔCTH^), or the CCM2^I428S^, CCM2^WD/AA^, and CCM2^I432D^ mutations all abolished CCM2 recruitment to mito-TLNRD1 (Fig. 5F and 5G). In addition, both the TLNRD1^2E^ and TLNRD1^2T^ double mutations abolished TLNRD1 recruitment to mito-CCM2. Altogether, our mutagenesis approach demonstrates that the TLNRD1-CCM2 interaction involves a hydrophobic groove on TLNRD1 and hydrophobic residues of CCM2.

### CCM2 modulates TLNRD1 localization, actin binding, and bundling activity

After identifying point mutations that disrupt the interaction between CCM2 and TLNRD1, we explored the cellular functions of the TLNRD1-CCM2 complex. We started by overexpressing the CCM2 mutants in endothelial cells. Surprisingly, deletion of CCM2^CTH^ (CCM2^ΔCTH^), or CCM2^WD/AA^, and CCM2^I432D^ mutations lead to a strong accumulation of CCM2 in the nucleus (Fig. S5A-B). This finding is particularly noteworthy because, although CCM2 is known to localize to the nucleus, the mechanisms controlling the nuclear translocation remain elusive (Swamy and Glading, 2022). A similar effect was seen with the patient mutation, CCM2^I428S^, suggesting it has a loss of function phenotype in cells. However, silencing TLNRD1 did not result in any noticeable changes in CCM2 subcellular localization (Fig. S5C-D), suggesting that the C-terminal helix in CCM2 likely serves functions beyond binding to TLNRD1. These findings complicate using these specific CCM2 mutants as tools for dissecting the role of the TLNRD1-CCM2 interaction. In addition, TLNRD1 silencing did not affect phospho-myosin light chain levels (Fig. S5E) and decreased rather than increased KLF4 expression in endothelial cells (Fig. S5F), both pathways regulated by CCM2 (Cuttano et al., 2016; Stockton et al., 2010). Therefore, it is likely that TLNRD1 acts downstream of CCM2 rather than directly regulating known CCM2 functions.

Next, we overexpressed our engineered TLNRD1 mutants in endothelial cells and conducted a detailed analysis of their subcellular localization (Fig. 6A). Intriguingly, both TLNRD1^2T^ and TLNRD1^2E^ demonstrated a subtle yet noticeable increase in nuclear localization when compared to TLNRD1^WT^ (Fig. 6B). Most remarkably, TLNRD1^2E^ failed to localize to actin stress fibers. At the same time, TLNRD1^2T^ exhibited a slightly enhanced localization to these structures compared to TLNRD1^WT^ (Fig. 6C).

**Fig. 6.**
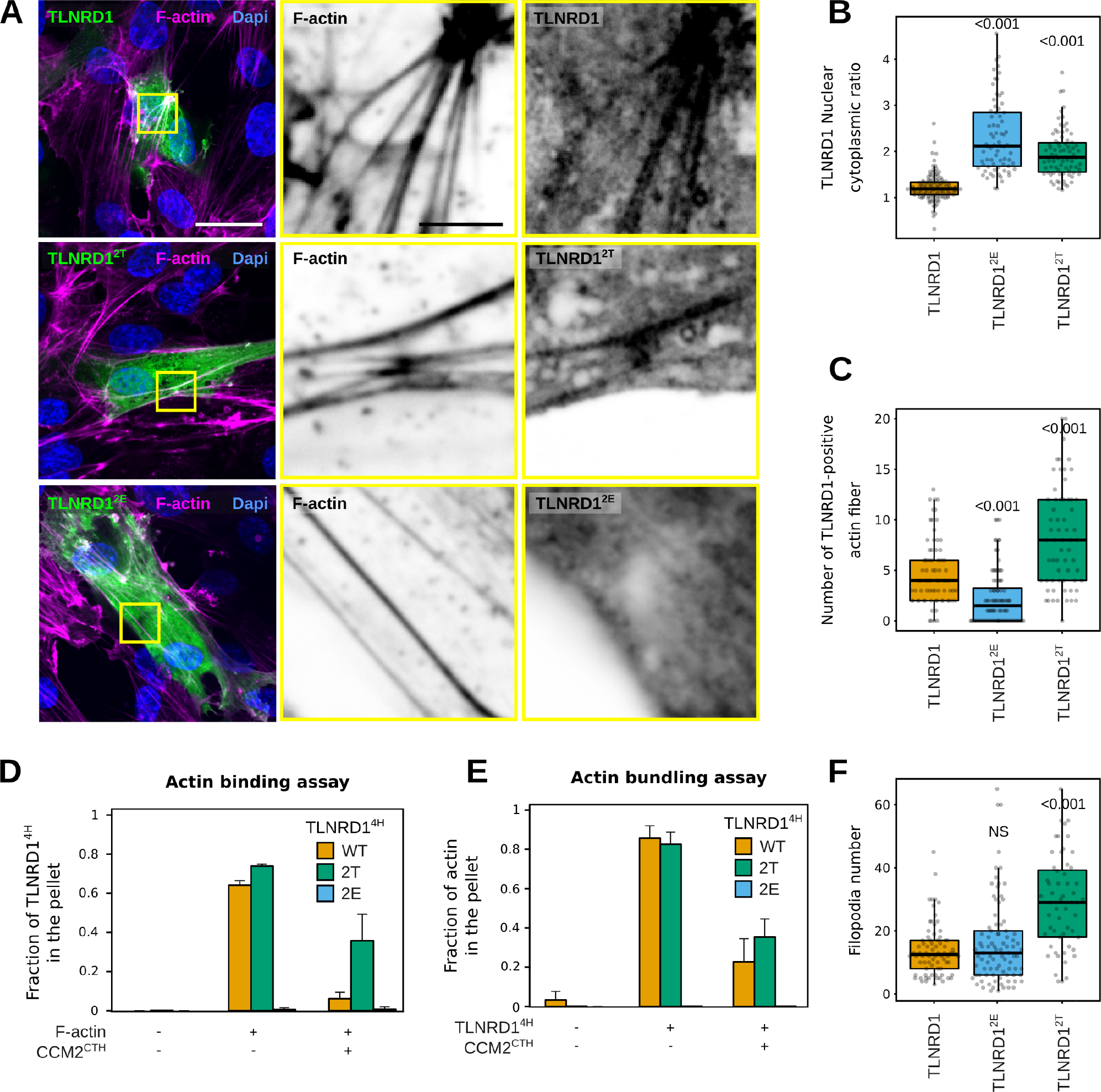
CCM2 modulates TLNRD1 localization and bundling activity. (**A**-**C**) HUVECs expressing various TLNRD1-GFP constructs were stained for DAPI and F-actin and imaged using a spinning disk confocal microscope. (**A**) Single Z-planes are displayed. The yellow squares highlight magnified ROIs. Scale bars: (main) 25 µm and (inset) 5 µm. (**B**-**C**) For each condition, the TLNRD1 nuclear-cytoplasmic ratio and the number of TLNRD1-positive actin fibers were quantified (see methods for details), and results are displayed as Tukey boxplots (three biological repeats, n> 72 cells per condition). (**D**-**E**) Actin co-sedimentation assay with various TLNRD1^4H^ mutants in the presence or absence of CCM2^CTH^. Centrifugation at high (**D**, 48,000 rpm) or low (**E**, 16,000 rpm) speeds can distinguish between F-actin binding and bundling capability. Quantifications of the gels shown in Fig. S6 are displayed. The quantification was performed using densitometry, and the fraction of TLNRD1 (**D**) and F-actin (**E**) present in the pellet was plotted. Standard deviation from three independent repeats are represented as error bars. (**F**) The number of filopodia in TLNRD1-positive cells described in (A) was manually quantified, and results are displayed as Tukey boxplots (three biological repeats, n> 72 cells per condition). For all panels, the P-values were determined using a randomization test. NS indicates no statistical difference between the mean values of the highlighted condition and the control.

These observations led us to hypothesize that the mutations we introduced in TLNRD1 could affect TLNRD1’s ability to bind to actin. Therefore, we carried out in vitro actin pulldown assays. These experiments showed that while both TLNRD1^WT^ and TLNRD1^2T^ could bind actin similarly, TLNRD1^2E^ could not. These findings concord with the hypothesis that the actin and CCM2 binding sites on TLNRD1 might overlap, rendering each TLNRD1 monomer incapable of binding to both actin and CCM2 simultaneously. To investigate this, we performed actin-binding assays in the presence or absence of CCM2^CTH^. These experiments revealed that TLNRD1^WT^ lost its ability to bind to actin in the presence of CCM2^CTH^ (Fig. 6D and Fig. S7A). Importantly, TLNRD1^2T^ maintained its ability to bind actin even in the presence of CCM2^CTH^, although the efficiency was somewhat reduced (Fig. 6D and Fig. S6A). Our results demonstrate that the TLNRD1-CCM2 and the TLNRD1-actin interactions are mutually exclusive.

To assess the functional relevance of this, we next investigated in vitro if CCM2^CTH^ would inhibit the ability of TLNRD1 to bundle actin. These experiments revealed that while TLNRD1^WT^ bundles actin effectively, as we reported previously (Cowell et al., 2021), the addition of CCM2^CTH^ inhibited TLNRD1^WT^ actin bundling. In contrast, TLNRD1^2E^ was unable to bundle actin. Given that TLNRD1 bundling activity contributes to filopodia formation in cancer cells (Cowell et al., 2021), we wanted to explore further the ability of our TLNRD1 mutants to modulate filopodia formation in HUVECs. The TLNRD1^2T^ mutant, which is unable to interact with CCM2, induced more filopodia than cells expressing TLNRD1^WT^ or TLNRD1^2E^ (Fig. 6F), providing further support that CCM2 inhibits TLNRD1 bundling activity and has functional relevance in cells.

## Discussion

Here, we report that TLNRD1 is primarily expressed in the vasculature in vivo and that silencing of TLNRD1 expression results in vascular phenotypes both in vitro and in vivo. We demonstrate that TLNRD1 is a member of the CCM complex and that TLNRD1 interacts directly and with high affinity to CCM2. We also show that the TLNRD1-CCM2 binding interface involves a hydrophobic groove on TLNRD1 and hydrophobic residues of CCM2. Altogether, we propose a model where TLNRD1 modulates endothelial cell functions by regulating actin organization. In our model, CCM2 binds to TLNRD1, sequestering TLNRD1 in the cytoplasm and inhibiting TLNRD1 bundling activity by competing with actin-binding. As TLNRD1 expression is altered in CCM lesions, we propose that TLNRD1 could be a novel actor in the CCM disease.

We find that TLNRD1 is a member of the CCM complex and that TLNRD1 depletion leads to vascular phenotypes both in vitro and in vivo. The CCM complex is best known for modulating and maintaining the vasculature, and loss of function mutations of CCM complex components lead to cerebral cavernous malformations (Fischer et al., 2013). Our results are consistent with work from others, as a recent preprint from Schnitzler et al. identified TLNRD1 to be linked to the CCM pathway (Schnitzler et al., 2022). Schnitzler et al. also reported that TLNRD1 targeting affects endothelial cell permeability and zebrafish heart development (Schnitzler et al., 2022). Interestingly, 75% of the familial CCM cases are attributed to mutations in KRIT, CCM2, and PDCD10 (Chohan et al., 2019). To our knowledge, TLNRD1 mutations have not yet been reported in CCM patients, and future work will aim at sequencing TLNRD1 in familial CCM samples. Instead, a reanalysis of previous work indicated that TLNRD1 expression is up-regulated in CCM lesions (Subhash et al., 2019). Interestingly, TLNRD1 expression is also up-regulated in KRIT1 but not in PDCD10 knock-out mice (Koskimäki et al., 2019b; a). TLNRD1 is also over-expressed in patients with dilated cardiomyopathy and ischemic heart disease (Liu et al., 2015) and is associated with significant stroke risk (Mishra et al., 2022). Taken together, TLNRD1 is emerging as an important regulator of endothelial cell function in vitro and in vivo that is misregulated in vascular diseases.

Mechanistically, it remains unclear how TLNRD1 regulates endothelial cell functions, but TLNRD1 bundling activity, as well as other TLNRD1 binding partners, are likely involved. Future work will aim to investigate the role of TLNRD1 in endothelial cells in more detail. Here, we found that TLNRD1 interacts with the CCM complex via CCM2. Our results are consistent with previous work as TLNRD1 was found to co-precipitate with CCM2 (Schnitzler et al., 2022), and TLNRD1 and CCM2 scored in a yeast two-hybrid screen (Luck et al., 2020). We identified that the TLNRD1-CCM2 interaction involves the C-terminal helix in CCM2 that interacts with TLNRD1’s 4-helix domain. Interestingly, this interaction is primarily hydrophobic, explaining why TLN1 R7R8 does not bind to CCM2 despite being structurally homologous to TLNRD1 (Cowell et al., 2021). Importantly, CCM2 binds to TLNRD1 on the same surface as actin binds, and we found that the TLNRD1-actin and TLNRD1-CCM2 interactions are mutually exclusive. Our results also indicate that CCM2 inhibits TLNRD1 actin-bundling activity. Interestingly, previous work reported that CCM2 deletion leads to an increase in actin stress fiber formation in endothelial cells (Faurobert et al., 2013), which may be consistent with our finding that CCM2 inhibits an actin-bundling protein. TLNRD1 interacts with CCM2 with high affinity. As the affinity of the TLNRD1-actin interaction is close to that of the TLNRD1-CCM2 interaction, an attractive hypothesis is that local actin dynamics could regulate the TLNRD1-CCM2 and the TLNRD1-actin interactions.

One key CCM complex function within cells is to restrain cellular contractility, thereby limiting the formation of stress fibers that compromise endothelial barrier function. Notably, each component of the CCM complex inhibits the RhoA-ROCK (Rho-associated coiled coil-forming kinase) pathway, subsequently preventing myosin light chain (MLC) phosphorylation (Riolo et al., 2021; Su and Calderwood, 2020; Stockton et al., 2010; Whitehead et al., 2009; Hartmann et al., 2015; Lisowska et al., 2018; Zheng et al., 2010; Crose et al., 2009). Yet, the intricate molecular dynamics underpinning how a deficiency in a CCM component triggers RhoA/ROCK activation is yet to be fully elucidated. Here, we found that CCM2 governs the cytoplasmic localization of TLNRD1, attenuating its ability to bundle actin. Given these insights, we posit a novel mechanism where the CCM complex can also influence actin cytoskeleton and vascular stability independently of RhoA, specifically by regulating TLNRD1’s bundling function.

## Material and methods

### Cells

U2OS osteosarcoma cells and HEK293 (human embryonic kidney) cells were grown in DMEM (Dulbecco’s Modified Eagle’s Medium; Sigma, D1152) supplemented with 10% fetal bovine serum (FCS) (Biowest, S1860). U2OS cells were purchased from DSMZ (Leibniz Institute DSMZ-German Collection of Microorganisms and Cell Cultures, Braunschweig DE, ACC 785). HEK293 cells were provided by ATCC (CRL-1573). Human Umbilical Vein Endothelial Cells (HUVEC) (PromoCell C-12203) were grown in Endothelial cell growth medium (ECGM) (PromoCell C-22010) supplemented with supplemental mix (Promocell C-39215) and 1% penicillin-streptomycin (Sigma). Endothelial primary cells from P0 (commercial vial) were expanded to a P3 stock stored in a -80°C freezer to standardize the experimental replicates.

### Antibodies and reagents

The anti-TLNRD1 antibody used for western blot (1:1000) was raised in rabbits against recombinantly expressed human TLNRD1 (residues 1–362) by Capra Science. The rabbit anti-TLNRD1 used to stain the brain section was purchased from Atlas Antibodies (1:200 for IF, HPA071716). Other rabbit antibodies used in this study include anti-KRIT1 (1:1000 for WB, Abcam, ab196025), anti-fibronectin (1:200 for IF, Sigma-Aldrich, f3648), anti-GFP (1:1000 for WB, LifeTechnologies, A11121) and anti-Myosin light chain (phospho S20) antibody (1:200 for IF, Abcam, ab2480). Mouse antibodies were anti-CCM2 (1:1000 for WB, Thermo Fisher Scientific, MA5-25668), anti-PECAM (1:200 for IF, Invitrogen, 37-0700), and anti-GAPDH (1:1000 for WB, Hytest, 5G4).

### Plasmids used for cell studies

The pMTS-mScarlet-I-N1 (mito-mScarlet) was a gift from Dorus Gadella (Addgene plasmid 85059; RRID: Addgene 85059) (Bindels et al., 2017). mScarlet-MYO10 was described previously and is available on Addgene (Addgene plasmid 145179; RRID: Addgene 145179) (Jacquemet et al., 2019). The GFP-TLNRD1 was as described previously (Gingras et al. 2010), and the GFP-TLNRD1-4H was generated from it. Both constructs are in the vector pEGFP-C1 and will be deposited in Addgene.

The ITG1BP1-GFP construct (ICAP-1) was a gift from Daniel Bouvard (University of Grenoble, FR). The PDCD10-mEmerald (PDCD10-GFP) and the CCM2-mCherry constructs were generated by the Genome Biology Unit core facility cloning service (Research Programs Unit, HiLIFE Helsinki Institute of Life Science, Faculty of Medicine, University of Helsinki, Biocenter Finland) by transferring the PDCD10 entry clone (100005213) into pcDNA6.2/N-emGFP-DEST and the CCM2 entry clone (100073308) into pcDNA6.2/C-mCherry-DEST using a standard LR reaction protocol. The mito-CCM2-mScarlet-I (mito-CCM2) and mito-PDCD10-mScarlet-I (mito-PDCD10) constructs were created by inserting a custom gene block (IDT) in the pMTS-mScarlet-I-N1 plasmid (Addgene 85059) using the XhoI/EcoRI sites. The gene blocks used are available in Table S2. The TLNRD1-5H-eGFP (TLNRD1^5H^), CCM2-wt-eGFP (CCM2WT), CCM2-d410-444-eGFP (CCM2ΔCTH), CCM2-I428S-eGFP (CCM2^I428S^), CCM2-I432D-eGFP (CCM2^I432D^), CCM2-W421AD422A-eGFP (CCM2^WD/AA^), TLNRD1-2T-eGFP (TLNRD1^2T^), TLNRD1-2E-eGFP (TLNRD1^2E^), mito-TLNRD1-mScarlet-I constructs were purchased from GenScript. Briefly, the gene fragments were synthesized using gene synthesis and cloned into pcDNA3.1(+)-N-eGFP using the BamHI/XhoI sites. The full plasmid sequences are available in Table S2. All constructs were sequence verified. These plasmids will be available on Addgene.

### Plasmids used for producing recombinant proteins

The FL and 4H TLNRD1 were described previously (Cowell et al., 2021) and are available on Addgene (Addgene plasmids 159384 and 159386). The FL CCM2 constructs were purchased from GeneArt and subcloned into pET151. GST-CCM2 constructs were produced by subcloning the CCM2 constructs into the XmaI/SacI sites of pET49b using ligation-independent cloning. The HisSUMO-tagged CCM2(413-438) was subcloned into a modified pET47b vector encoding a hexahistidine tag fused to SUMO (where all lysine residues were mutated to arginine residues) followed by an HRV-3C cleavage site (provided by Dr. I.A. Taylor, The Francis Crick Institute, UK) using ligation-independent cloning. Mutations were introduced by site-directed mutagenesis. All constructs were sequence verified. The recombinant expression vectors of GST-CCM2(FL), HisSumo-CCM2(413-438), TLNRD1(4H)-2T, TLNRD1(4H)-2E will be deposited in Addgene.

### Cell transfection

U2OS cells were transfected using Lipofectamine 3000 and the P3000™ Enhancer Reagent (Thermo Fisher Scientific, L3000001) according to the manufacturer’s instructions. HUVECs were transfected using the Neon™ Transfection System (Thermo Fisher Scientific, MPK5000) and the Neon™ Transfection System 10 µL Kit (Thermo Fisher Scientific, MPK1025) according to the manufacturer’s instructions. Briefly, 150k cells and 1 µg of plasmid DNA were used for each transfection (1.350 pulse voltage and 30 ms pulse width). Transfected cells were seeded in a glass bottom µ-slide 8 well (Ibidi, 80807) with a glass bottom precoated with warm ECGM without antibiotics. 50k untransfected cells/well were added after transfection. Cells were then grown for 48 h and fixed using pre-warmed paraformaldehyde 4% in PBS (Thermo Fisher Scientific, 28908) for 10 min at 37°C. HEK293T cells were transfected using 100x polyethyleneimine reagent (Sigma-Aldrich). Cells were seeded in a 15 cm dish and allowed to reach 80% confluence before transfection. The transfection mixture consisted of 8 µg plasmids diluted in 500 ml OptiMEM and 54 µl of 1x polyethyleneimine (diluted with 150 mM NaCl) pre-incubated with 446 ml of OptiMEM for 5 min at room temperature. Transfection mixtures were incubated for 20 min at room temperature. The medium was then removed from the cells and replaced with 10 ml growth medium and the transfection mixture. Following 10 h of incubation, the transfection mixture was removed, a fresh medium was added, and transfected cells were used the following day.

### siRNA-mediated gene silencing

The expression of KRIT1 was suppressed in U2OS cells using 83 nM siRNA and Lipo-fectamine™ 3000 (Thermo Fisher Scientific, L3000001) according to the manufacturer’s instructions. The expression of TLNRD1 was suppressed in HUVECs using 50 nM siRNA and Lipofectamine™ RNAiMAX (Thermo Fisher Scientific, 13778075) following the manufacturer’s instructions. HUVECs were seeded on fibronectin-coated wells or microchannels in the siRNA-containing solution for 2 h. Then, normal media was added on top of the cells. The following day, the transfection media was replaced. These additional steps maximized siRNA entry in the cells while preserving cell viability. In all cases, cell experiments were performed after 72 h of treatment with siRNA. siRNAs used were AllStars Negative siRNA (SI03650318), TLNRD1 siRNA 1 (SI04314569) TLNRD1 siRNA 2 (SI04362820), KRIT1 siRNA 1 (SI02777173), KRIT1 siRNA 2 (SI03054499), all from Qiagen.

### Primers and qPCR

RNAs were extracted and purified using an RNeasy Mini Kit (Qiagen, 74104) according to the manufacturer’s instructions, including a DNase digestion using an RNase-Free DNase (Qiagen, 79254). The RNA concentration and quality were assessed using a Nanodrop 2000 (Ther-mofisher). cDNAs were then synthesized using iScript cDNA (bio-rad, 1708890) according to the manufacturer’s instructions. Real-time quantitative PCR (RT-qPCR) was performed using QuantStudio 3 (Thermofisher, A28567) with PowerUp SYBR Green Mastermix (Life Technologies, A25741). The reaction mixture consisted of 4 ng of cDNA, primers, and master mix and was run under the following conditions: Hold (50°C for 2 minutes, 95°C for 2 minutes), PCR (95°C for 15 seconds, 60°C for 1 minute) and Melt Curve (95°C for 15 seconds, 60°C for 1 minute and 95°C for 15 seconds). The expression levels of the target genes were normalized to the expression level of GAPDH using the [ΔΔCt method]. All experiments were performed in triplicate, and the results were analyzed using QuantStudio Design Analysis Software 2.6.0.

### SDS-PAGE and quantitative western blotting

Protein extracts were separated under denaturing conditions by SDS-PAGE and transferred to nitrocellulose membranes using a Mini Blot Module (Invitrogen, B1000). Membranes were blocked for 30 minutes at room temperature using 1× Start-ingBlock buffer (Thermo Fisher Scientific, 37578). After blocking, membranes were incubated overnight with the appropriate primary antibody (1:1000 in blocking buffer), washed three times in PBS, and probed for 1 h using a fluorophore-conjugated secondary antibody diluted 1:5000 in the blocking buffer. Membranes were washed thrice using PBS over 30 min and scanned using an iBright FL1500 imaging system (Invitrogen).

### GFP-trap pulldown

Cells transiently expressing bait GFP-tagged proteins were lysed in a buffer containing 20 mM HEPES, 75 mM NaCl, 2 mM EDTA, 1% NP-40, as well as a cOmplete™ protease inhibitor tablet (Roche, 5056489001), and a phosphatase inhibitor mix (Roche, 04906837001). Lysates were then centrifuged at 13,000 g for 10 min at 4C. Clarified lysates were incubated with GFP-Trap agarose beads (Chromotek, gta-20) overnight at 4C. Complexes bound to the beads were isolated by centrifugation, washed three times with ice-cold lysis buffer, and eluted in Laemmli reducing sample buffer.

### Mass spectrometry analysis

Affinity-captured proteins were separated by SDS-PAGE and allowed to migrate 10 mm into a 4-12% polyacrylamide gel. Following staining with InstantBlue (Expedeon), gel lanes were sliced into five 2-mm bands. The slices were washed using a solution of 50% 100 mM ammonium bicarbonate and 50% acetonitrile until all blue colors vanished. Gel slices were washed with 100% acetonitrile for 5-10 min and then rehydrated in a reducing buffer containing 20 mM dithiothreitol in 100 mM ammonium bicarbonate for 30 min at 56°C. Proteins in gel pieces were then alkylated by washing the slices with 100% acetonitrile for 5-10 min and rehydrated using an alkylating buffer of 55 mM iodoacetamide in 100 mM ammonium bicarbonate solution (covered from light, 20 min). Finally, gel pieces were washed with 100% acetonitrile, followed by washes with 100 µl 100 mM ammonium bicarbonate, after which slices were dehydrated using 100% acetonitrile and fully dried using a vacuum centrifuge. Trypsin (0.01 µg/µl) was used to digest the proteins (37°C overnight). After trypsinization, an equal amount of 100% acetonitrile was added, and gel pieces were further incubated at 37°C for 15 min, followed by peptide extraction using a buffer of 50% acetonitrile and 5% formic acid. The buffer with peptides was collected, and the sample was dried using a vacuum centrifuge. Dried peptides were stored at -20°C. Before LC-ESI-MS/MS analysis, dried peptides were dissolved in 0.1% formic acid. The LC-ESI-MS/MS analyses were performed on a nanoflow HPLC system (Easy-nLC1000, Thermo Fisher Scientific) coupled to the Q Exactive mass spectrometer (Thermo Fisher Scientific, Bremen, Germany) equipped with a nano-electrospray ionization source. Peptides were first loaded on a trapping column and subsequently separated inline on a 15 cm C18 column (75 µm × 15 cm, ReproSil-Pur 5 µm 200 Å C18-AQ, Dr. Maisch HPLC GmbH, Ammerbuch-Entringen, Germany). The mobile phase consisted of water with 0.1% formic acid (solvent A) and acetonitrile/water (80:20 (v/v)) with 0.1% formic acid (solvent B). A linear 30 min gradient from 6% to 39% was used to elute peptides. MS data was acquired automatically by using Thermo Xcalibur 3.0 software (Thermo Fisher Scientific). An information-dependent acquisition method consisted of an Orbitrap MS survey scan of mass range 300-2000 m/z followed by HCD fragmentation for the 10 most intense peptide ions. Raw data from the mass spectrometer were submitted to the Mascot search engine using Proteome Discoverer 1.5 (Thermo Fisher Scientific). The search was performed against the human database SwissProt-2018-04, assuming the digestion enzyme trypsin, a maximum of two missed cleavages, an initial mass tolerance of 10 ppm (parts per million) for precursor ions, and a fragment ion mass tolerance of 0.020 Dalton. Cysteine carbamidomethylation was set as a fixed modification, and methionine oxidation was set as a variable modification. To generate the TLNRD1 dataset, three biological replicates were combined. Proteins enriched at least threefold in TLNRD1-GFP over GFP (based on normalized spectral count) and detected with at least six spectral counts (across all repeats) were considered putative TLNRD1 binders. The fold-change enrichment and the significance of the association used to generate the volcano Plot (Fig. 1B) were calculated in Scaffold 5 (version 5.2.0; Proteome Software Inc.) using a Fisher’s exact t-test corrected using the Benjamini-Hochberg method. To generate the protein-protein interaction network displayed in (Fig. 1C), enriched proteins were mapped onto a merged human interactome consisting of PPIs reported in STRING (v. 11.5) (Szklarczyk et al., 2019) directly in Cytoscape (version 3.9.1) (Shannon et al., 2003).

### Zebrafish experiments

The adult zebrafish of kdrl:mCherry-CAAX(s916) strain (Hogan et al., 2009) were housed in an Aqua Schwarz stand-alone rack (Aqua Schwarz GmbH). The husbandry of adult fish was carried out under license no. MMM/465/712-93 (issued by the Finnish Ministry of Forestry and Agriculture). Embryos were obtained via natural spawning in breeding tanks. Single-guide RNAs (sgRNA) targeting the tlnrd1 locus were designed using CHOPCOP software (Labun et al., 2019). SgRNAs were synthesized using a sgRNA synthesis kit (New England Biolabs) as described by the manufacturer and purified using RNA-clean 25 columns (ZYMO Research) and using the following oligonucleotides designed to target tlnrd1: tlnrd1-sgRNA9 (5’-TTC TAA TAC GAC TCA CTA TAG CTC GGG GAA ATC AGA TAG CGG TTT TAG AGC TAG A-3’), tlnrd1-sgRNA14 (5’-TTC TAA TAC GAC TCA CTA TAG TTA GCG GCA GCT TGC AAC AAG TTT TAG AGC TAG A-3’), tlnrd1-sgRNA24 (5’-TTC TAA TAC GAC TCA CTA TAG CTA TGG CTA GTA GTG GCT CGG TTT TAG AGC TAG A-3’) and tlnrd1-sgRNA25(5’-TTC TAA TAC GAC TCA CTA TAG CAC ACT ACT ATG GCT AGT AGG TTT TAG AGC TAG A-3’). As a control, sgRNAs targeting the slc45a2 locus were used (Heliste et al., 2020). Purified sgRNAs targeting tlnrd1 and slc45a2 were pooled separately and complexed with recombinant Cas9 protein (New England Biolabs) in vitro using 300mM KCl buffer for 5 minutes at +37°C. The solution was injected into 1-4-cell stage zebrafish embryos. The efficacy of tlnrd1 targeting was confirmed by extracting DNA as described earlier (Meeker et al., 2007) and amplifying targeted regions using nested-PCR (tlnrd1-L3 (5’-TCAT TTA CAT GGC ACG AAG AAC-3’), tlnrd1-R4 (5’-GGT GAG GTT CTT CAG GAT GTT C-3’), tlnrd1-L4 (5’-CGA GTG AAG TTT CAT GTT TTC G-3’), tlnrd1-R3 (5’-ATA GAC AGC TCC TTG GTT CTG G-3’)) and Sanger sequencing of amplicons. TIDE software (Brinkman et al., 2014) was used to determine the mutagenesis efficacy from sequencing chromatograms. Zebrafish embryos were incubated at 28.5C in E3 medium (5 mM NaCl, 0.17 mM KCl, 0.33 mM CaCl_2_, 0.33 mM MgSO_4_) supplemented with 0.2 mM phenyl-thiourea until anesthetized and analyzed. Embryos were imaged using a Nikon Eclipse Ti2 widefield microscope equipped with a 2x objective or a Zeiss stereoLumar fluorescence stereomicroscope. Vascular measurements were manually measured using Fiji. Heart rate measurements were conducted using Fiji using kymographs (Schindelin et al., 2012).

### Light microscopy setup

The spinning-disk confocal microscope used was a Marianas spinning-disk imaging system with a Yokogawa CSU-W1 scanning unit on an inverted Zeiss Axio Observer Z1 microscope controlled by SlideBook 6 (Intelligent Imaging Innovations, Inc.). Images were acquired using either an Orca Flash 4 sCMOS camera (chip size 2,048 × 2,048; Hamamatsu Photonics) or an Evolve 512 EMCCD camera (chip size 512 × 512; Photometrics). The objectives used were 40x (NA 1.1 water, Zeiss LD C-Apochromat) and 63x oil (NA 1.4 oil, Plan-Apochromat, M27) objectives. The structured illumination microscope (SIM) used was DeltaVision OMX v4 (GE Healthcare Life Sciences) fitted with a 60x Plan-Apochromat objective lens, 1.42 NA (immersion oil RI of 1.516) used in SIM illumination mode (five phases x three rotations). Emitted light was collected on a front-illuminated pco.edge sCMOS (pixel size 6.5 mm, readout speed 95 MHz; PCO AG) controlled by SoftWorx. The confocal microscope used was a laser scanning confocal microscope LSM880 (Zeiss) equipped with an Airyscan detector (Carl Zeiss) and a 40x water (NA 1.2) or 63x oil (NA 1.4) objective. The microscope was controlled using Zen Black (2.3), and the Airyscan was used in standard super-resolution mode.

### Quantification of endothelial monolayer integrity

We used a fibronectin accessibility assay to measure the integrity of the monolayer formed by HUVEC upon TLNRD1 silencing. SiRNA treatment was done while HUVECs were attached to glass-bottomed 24-well plates or microchannels (Ibidi µ-slide I LUER 0.4, 80177) previously coated with 10 µg/ml of fibronectin (Sigma Aldrich, 341631). In the flow-stimulated samples, the perfusion of the microchannels was done for 24 h (starting at 48 h post-siRNA treatment) with a perfusion speed of 400 µm/sec (Follain et al., 2018; Osmani et al., 2021). At 72 h post-siRNA treatment, cells were fixed using pre-warmed paraformaldehyde 4% in PBS (Thermo Fisher Scientific, 28908) for 10 min at 37°C. When indicated, samples were stained directly so only the fibronectin localized underneath antibody-permeable junctions would be labeled. When samples were permeabilized to visualize all the deposited fibronectin, cells were incubated with a solution of 0.2% Triton X-100 in PBS (Sigma Aldrich, 9002-93-1) for 10 min at room temperature. Samples were then incubated with the primary antibodies diluted in PBS (1:200) for 1 h at room temperature. After three PBS washes, samples were incubated in the dark with the fluorescently conjugated secondary antibodies diluted in PBS (1:400) for 30 min at room temperature. Samples were imaged in 3D using a spinning disk confocal microscope, and the images were analyzed using Fiji. Briefly, after generating maximal intensity projections, the FN patches were automatically segmented using the “default” thresholding method and the “run analyze particles” function. The percentage of the field of view covered by fibronectin patches was then measured.

### Brain slice preparation

All animal experiments were ethically assessed and authorized by the National Animal Experiment Board and following The Finnish Act on Animal Experimentation (Animal license number ESAVI/12558/2021). Mice (females Hsd: Athymic Nude-Foxn/1nu, Envigo) were housed in standard conditions (12-h light/dark cycle) with food and water available ad libitum. To collect brains, mice were sacrificed and placed in a ventral-down position. Incision at the base of the skull and cutting on both sides toward the sphenoid allowed the extraction of the brain in a single piece. Brains were rinsed in PBS and fixed in paraformaldehyde 4% (Thermo Fisher Scientific, 043368.9M) at 4° overnight.

Extracted brains were embedded in low melting point agarose (Thermo Fisher Scientific, 16520050), and 100 to 200 µm thick brain slices were prepared using a vibratome (Leica VT1200 S). Brain slices were stored in a 24-well plate and stained using a protocol adapted from (Fercoq et al., 2020). Briefly, sections were incubated with the permeabilization buffer (in PBS: 10% Horse Serum (v/v, Gibco, 16050-122), 1% BSA (w/v, Sigma-Aldrich, 1003435812), 0.3% TX-100 (v/v, Sigma-Aldrich, 9002-93-1) and 0.05% Sodium azide (w/v, Sigma-Aldrich, 26628-22-8) for 1 h at room temperature before being rinsed once with the washing buffer (in PBS: 1% BSA, 0.1% TX-100 (v/v) and 0.05% Sodium azide (w/v). Samples were then incubated with primary antibodies, diluted in the antibody dilution buffer (In PBS: 10% Horse Serum (v/v), 1% BSA (w/v), 0.1% TX-100 (v/v) and 0.05% Sodium azide (w/v), on a rocker for 3 h at RT. Samples were washed thrice for 20 min with the washing buffer while on a rocker. Next, fluorescently conjugated secondary antibodies, diluted in the antibody dilution buffer, were added to the wells, and the plate was placed on a rocker for 3 h at RT. Samples were washed twice for 5 min with the washing buffer while on a rocker. Finally, the samples were rinsed for 5 min with PBS and then mounted on a slide using ProLong™ Glass Antifade Mountant (Thermo Fisher Scientific, P36980). Brain slices were then imaged using a spinning disk confocal microscope.

### Nuclear to cytoplasmic ratio measurements

HUVECs expressing the indicated CCM2 or TLNRD1 constructs were allowed to form a monolayer for three days before being fixed and stained for PECAM and Dapi. High-resolution imaging of the monolayers was conducted using a spinning disk confocal microscope. Subsequently, to quantify the nuclear-to-cytoplasmic ratio of TLNRD1 and CCM2, SUM projections of the acquired image stacks were generated using Fiji. Automatic cell and nuclei segmentation was performed using the ZeroCostDL4Mic cellpose notebook (Pachitariu and Stringer, 2022; von Chamier et al., 2021). Two separate models within Cellpose were employed—the ‘cyto2’ model for segmenting the transfected cells and the ‘nuclei’ model for segmenting the nuclei. Each segmentation was then manually validated within Fiji, following which cytoplasmic segmentation masks were created by excluding the nuclear mask from the overall cell mask. Finally, the average integrated fluorescence density within the nucleus was calculated and normalized against that in the cytoplasm to obtain the desired ratio.

### Mitochondrial trapping experiments

Cells expressing the constructs of interest were plated on fibronectin-coated glass-bottom dishes (MatTek Corporation) for 2 h. Samples were fixed for 10 min using a solution of 4% PFA, then permeabilized using a solution of 0.25% (v/v) Triton X-100 for 3 min. Cells were then washed with PBS and quenched using a solution of 1 M glycine for 30 min. Samples were then washed three times in PBS and stored in PBS containing SiR-actin (100 nM; Cytoskeleton; CY-SC001) at 4°C until imaging. Just before imaging, samples were washed three times in PBS. Images were acquired using a spinning-disk confocal microscope (100× objective). The Pearson correlation coefficients were calculated from the 3D stacks using the JaCoP Fiji plugin (Schindelin et al., 2012; Bolte and Cordelières, 2006).

### Protein expression and purification

BL21(DE3)* competent cells were transformed with the relevant plasmid and grown in lysogeny broth supplemented with appropriate antibiotic (TLNRD1: 100 µg/mL ampicillin; CCM2: 50 µg/mL kanamycin) until the OD600 reached 0.6. Protein expression was induced by adding 0.4 mM IPTG and expressed overnight at 20°C. Cells were harvested by centrifugation, resuspended in lysis buffer (50 mM Tris-HCl pH 8, 250 mM NaCl, 5% v/v glycerol) at 5 mL per gram of cells, and stored at -80°C.

TLNRD1 and SUMO-CCM2(413-438) proteins were purified using nickel-affinity chromatography and by ion-exchange chromatography. Briefly, cells were thawed, supplemented with 1 mM PMSF, 5 mM DTT, 0.2% v/v Triton X-100, and lysed by sonication. Cell debris were removed by centrifugation, and the supernatant was filtered and loaded onto a 5 mL HisTrap HP column (Cytiva) using an AKTA Start (GE Healthcare). The column was washed in buffer containing 50 mM Tris-HCl (pH 8), 600 mM NaCl, 30 mM imidazole, 4 mM MgCl2, 4 mM ATP, 5% v/v glycerol, 0.2% v/v Triton X-100, 5 mM DTT. Bound protein was eluted using a 75 mL linear gradient of 0 - 300 mM imidazole. Fractions containing the protein of interest were pooled and diluted with 5 volumes of either 20 mM sodium phosphate (pH 6.5, TLNRD1) or 20 mM Tris-HCl (pH 8, SUMO-CCM2(413-438)) and loaded onto either a 5 mL HiTrap SP (TLNRD1) or a 5 mL HiTrap Q (SUMO-CCM2) (Cytiva) and eluted with a 75 mL linear gradient of 0 - 750 mM NaCl. Purified proteins were dialyzed overnight against PBS (pH 7.4) supplemented with 5 mM DTT, snap-frozen in LN_2_, and stored at -80°C.

### GST pulldown assay

Cell pellets of the various GST-CCM2 constructs were thawed and lysed as above. The filtered supernatant was batch-bound to Glutathione Superflow Agarose resin (Thermo Fisher Scientific) for 90 min whilst being rolled at 4°C. Unbound proteins were removed by centrifugation/aspiration, and the resin was washed five times with 10 volumes of PBS (pH 7.4). 50 µL washed resin containing immobilized GST-CCM2 was incubated with 200 µL purified protein at 40 µM for 60 min at room temperature with inversion mixing. The unbound protein was removed by centrifugation/aspiration. The resin was subjected to 5 × 500 µL washes of PBS (pH 7.4), resuspended in 50 µL PBS (pH 7.4), and analyzed by SDS-PAGE.

### Molecular modelling

To produce the structural models of the TLNRD1-CCM2 complex, the sequence of TLNRD1(FL) and CCM2(410-444) were submitted to the protein structure modeling tool, ColabFold (Mirdita et al., 2022) and the crystal structure of TLNRD1 (PDB accession code 6XZ4) was used as a template. All figures were made using PyMOL (Version 2.5; Schrödinger, LLC).

### Fluorescence Polarization assay

The SUMO-CCM2(413-438) contains a single cysteine (C437) that was used to couple the fusion protein with a maleimide-fluorescein dye (Thermo Fisher Scientific) following the manufacturer’s protocol. Assays were performed in triplicate with a 2-fold serial protein dilution, with target peptides at 500 nM. Fluorescence polarization was measured using a CLARIOstar plate reader (BMGLabTech) at 25°C (excitation: 482 ± 8 nm; emission: 530 ± 20 nm). Data were analyzed using GraphPad Prism 8 software, and Kd values were generated using the one-site total binding equation.

### Actin preparation

Rabbit skeletal muscle acetone powder was kindly gifted by Professor Mike Geeves (University of Kent, UK), and actin was prepared using cycles of polymerization/depolymerization following the protocol of Spudich and Watt (Spudich and Watt, 1971). Briefly, 1 g of acetone powder was stirred on ice in extraction buffer (10 mM Tris pH 8, 0.5 mM ATP, 0.2 mM CaCl_2_, 1 mM DTT), filtered and clarified by ultracentrifugation (30,000 rpm, 70Ti rotor, 1 h). The supernatant was supplemented with 3 M KCl and 1 M MgCl_2_ to final concentrations of 100 mM and 2 mM, respectively, stirred at room temperature for 1 h, and centrifuged for 3 h at 30,000 rpm. The pellet of F-actin was resuspended in a depolymerization buffer (5 mM Tris pH 7.5, 0.2 mM CalCl_2_, 1 mM NaN_3_), gently homogenized, and dialyzed against depolymerization buffer overnight. The dialyzed G-actin was centrifuged (30,000 rpm, 1 h), and the supernatant was retained, supplemented with 0.2 mM ATP and 3% w/v sucrose, and snap-frozen in LN_2_.

### F-actin co-sedimentation assays

G-actin was polymerized by adding ATP, KCl, and MgCl2 to final concentrations of 5 µM, 100 mM, and 2 mM, respectively. The polymerized actin was recovered by centrifugation (30,000 rpm, 1 h), gently homogenized in co-sedimentation buffer (10 mM Tris pH 7.5, 100 mM NaCl, 2 mM MgCl_2_, 1 mM DTT, 1 mM NaN_3_) and stored at 4°C. Co-sedimentation assays were performed with F-actin at 15 µM and equimolar concentrations of the other protein components. Binding was carried out at room temperature for 1 h, and the reaction was centrifuged at either 16,000 rpm (bundling assay) or 48,000 rpm (binding assay) in a TLA-100 ultracentrifuge rotor for 20 minutes. Equal volumes of supernatant and pellet fractions were analyzed by SDS-PAGE. Gels were quantified using ImageJ (Schneider et al., 2012)

### Quantification and statistical analysis

Randomization tests were performed using the online tool PlotsOfDifferences (Goedhart, 2019). Dot plots were generated using PlotsOfData (Postma and Goedhart, 2019). Volcano Plots were generated using VolcaNoseR (Goedhart and Luijster-burg, 2020).

### Data availability

Plasmids generated in this study are being deposited in Addgene. The mass spectrometry proteomics data have been deposited to the ProteomeXchange Consortium via the PRIDE (Perez-Riverol et al., 2022) partner repository with the dataset identifier PXD045258. The raw microscopy images used to make the figures are available on Zenodo (10.5281/zenodo.8377287). The authors declare that the data supporting the findings of this study are available within the article and from the authors upon request. Any additional information required to reanalyze the data reported in this paper is available from the corresponding authors.

## Acknowledgements

This study was supported by the Research Council of Finland (338537 to G.J.; 325464 to J.I.; and 332402 to G.F.), the Sigrid Juselius Foundation (to G.J. and to J.I.), the Cancer Society of Finland (Syöpäjärjestöt; to G.J. and to J.I.), and the Solutions for Health strategic funding to Åbo Akademi University (to G.J.). This research was supported by the InFLAMES Flagship Programme of the Academy of Finland (decision numbers: 337530, 337531, 357910, and 357911). This work was also supported by the Finnish Cancer Institute (K. Albin Johansson Professorship, J.I.), the Centre of Excellence program (346131, J.I.), and the Jane and Aatos Erkko Foundation (J.I.). B.T.G. was supported by the Biotechnology and Biological Sciences Research Council grant (BB/S007245/1) and the Cancer Research UK Program Grant (CRUK-A21671). The Cell Imaging and Cytometry Core facility, the Zebrafish Core (both at Turku Bioscience, University of Turku, Åbo Akademi University, and supported by Biocenter Finland), Turku Bioimaging and the Genome Biology Unit (Research Programs Unit, HiLIFE Helsinki Institute of Life Science, Faculty of Medicine, University of Helsinki, Biocenter Finland) are acknowledged for services, instrumentation, and expertise. Mass spectrometry was performed at the Turku Proteomics Facility, University of Turku and Åbo Akademi University. Biocenter Finland supports the facility.

## AUTHOR CONTRIBUTIONS

Conceptualization, G.J., and B.T.G.; Methodology, N.J.B., S.G., G.F., I.P., B.T.G., and G.J.; Formal Analysis, N.J.B., S.G., G.F., A.O.P., I.P., B.T.G., and G.J.; Investigation, N.J.B., S.G., G.F., A.O.P., M.V., A.R.C., B.B., I.P., B.T.G., and G.J.; Writing – Original Draft, B.T.G. and G.J.; Writing – Review and Editing, Everyone; Visualization, N.J.B., B.T.G., G.F. and G.J.; Supervision, I.P., B.T.G. and G.J.; Funding Acquisition, B.T.G. and G.J.

GJ is responsible for the experiments performed in the cells and tissue section. BTG is responsible for the experiments performed with recombinant proteins. IP is responsible for the Zebrafish experiments.

---

**Fig. S1.**
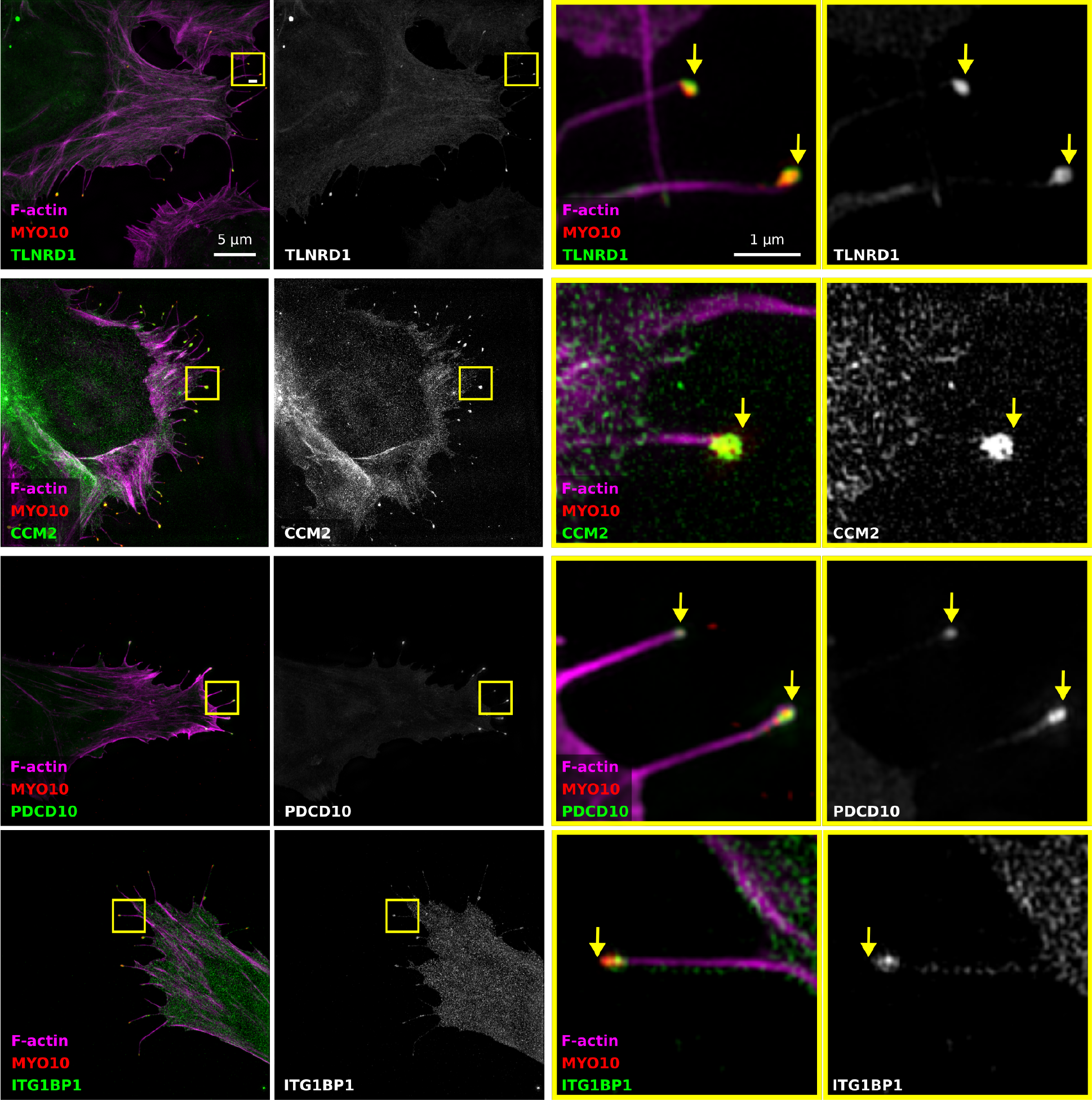
TLNRD1, CCM2, PDCD10, and ITG1BP1 localize at the tip of MYO10 filopodia. U2OS cells expressing mScarlet-MYO10 with TLNRD1-GFP, CCM2-GFP, PDCD10-GFP, or ITG1BP1-GFP were plated on fibronectin for 2 h, fixed and stained to visualize Factin. Samples were imaged using structured illumination microscopy. Representative maximum intensity projections are displayed; scale bars: (main) 5 µm; (inset) 1 µm. The yellow squares highlight magnified ROIs. The yellow arrows indicate the filopodia tips.

**Fig. S2.**
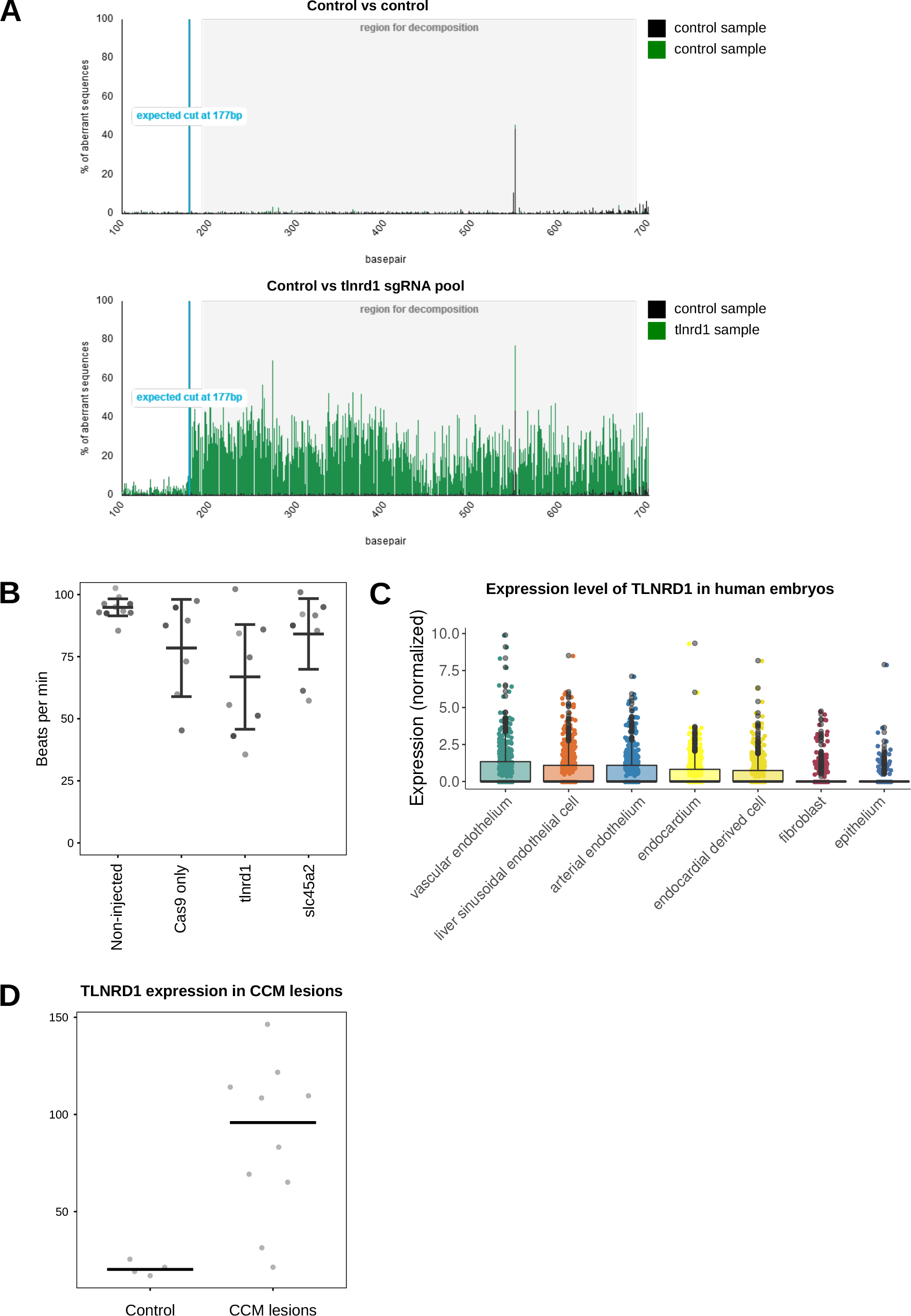
TLNRD1 in vivo. (**A**) Efficacy analysis of tlnrd1 CRISPR in zebrafish embryos. The Sanger sequencing chromatograms between control and tlnrd1 sgRNA injected samples were compared using TIDE software. The peak intensities show deviation of sequences and indicate effective editing of the tlnrd1 locus. (**B**) Zebrafish heart rate analysis. Zebrafish embryos were imaged using fast video microscopy, and heart rate was analyzed using kymographs in Fiji. (**C**) TLNRD1 expression in human embryos in single cells in various endothelial compartments, fibroblasts, and epithelial cells (data from (Xu et al., 2023)). (**D**) TLNRD1 expression in CCM lesions. Data from (Subhash et al., 2019). Controls are four patients diagnosed with temporal lobe epilepsy.

**Fig. S3.**
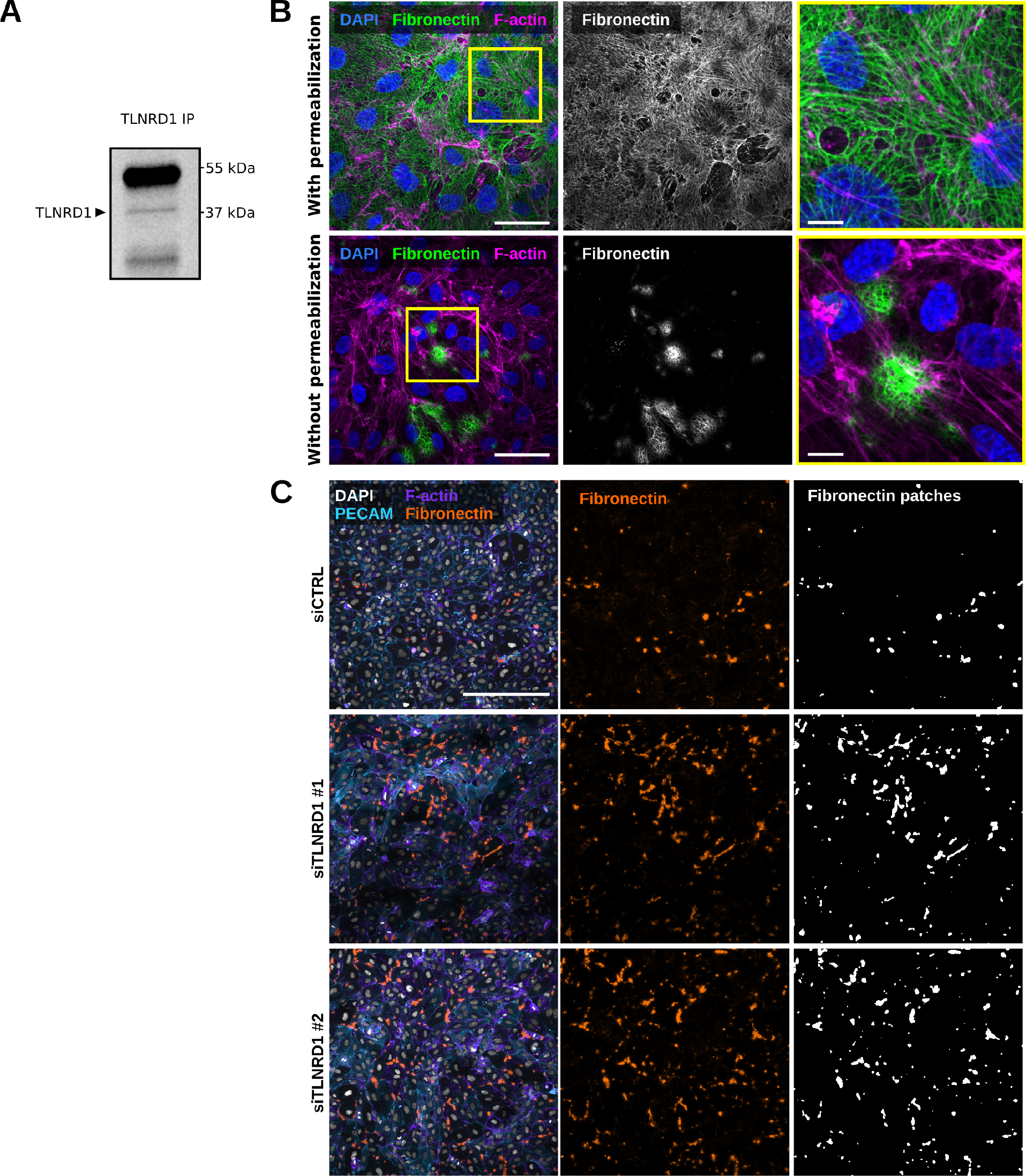
TLNRD1 in endothelial cells. (**A**) TLNRD1 immunoprecipitation from HUVEC lysate. A representative western blot is displayed. (**B**) HUVECs were allowed to form a monolayer. Cells were then fixed and stained for DAPI, F-actin, and Fibronectin (with or without permeabilization) before being imaged on a spinning disk confocal microscope. Representative maximum intensity projections are displayed. Scale bars: (main) 25 µm and (inset) 5 µm. (**C**) TLNRD1 expression was silenced in HUVECs using two independent siRNA. HUVECs were then allowed to form a monolayer without flow stimulation. Cells were then fixed and stained for DAPI, F-actin, PECAM, and Fibronectin (without permeabilization) before being imaged on a spinning disk confocal microscope. Representative maximum intensity projections are displayed. Scale bar: 250 µm.

**Fig. S4.**
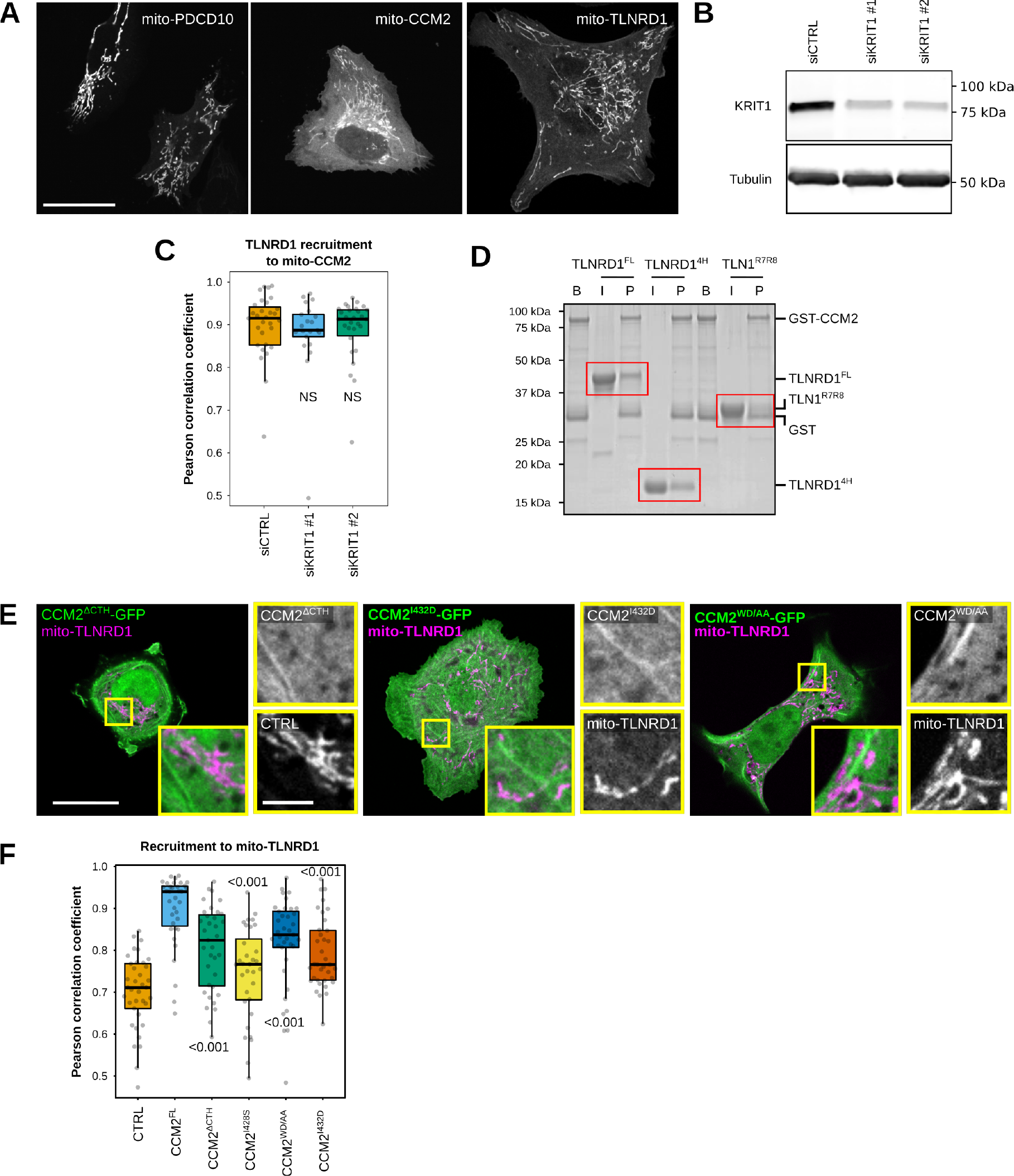
TLNRD1 interacts with CCM2. (**A**) U2OS cells expressing mito-PDCD10-mScarlet, mito-CCM2-mScarlet, or mito-TLNRD1-mScarlet were imaged using a spinning disk confocal microscope. (**B**) U2OS cells treated with siRNA targeting KRIT1 or siRNA control. KRIT1 levels were then analyzed using western blots. A representative western blot is displayed. (**C**) U2OS cells treated with siRNA targeting KRIT1 or siRNA control and expressing TLNRD1-GFP and mito-CCM2-mScarlet were imaged using a spinning disk confocal microscope. 3D colocalization analyses were performed using the JACoP Fiji plugin, and results are displayed as Tukey boxplots (three biological repeats, n> 21 image stacks per condition). (**D**) Glutathione agarose-bound GST-CCM2 (beads: B) was incubated with recombinant TLNRD1, TLNRD1^4H^, or TLN1^R7R8^ (input: I). After multiple washes, proteins bound to the beads (pellet: P) were eluted. The various fractions were then analyzed using SDS-PAGE followed by coomassie staining. A representative gel of 3 independent repeats is displayed. Red boxes highlight areas of interest in the gel. (**E**) U2OS cells expressing various GFP-tagged CCM2 constructs and mito-TLNRD1-mScarlet or mito-mScarlet (CTRL) were imaged using a spinning disk confocal microscope. Representative single Z-planes are displayed. The yellow squares highlight magnified ROIs. Scale bars: (main) 25 µm and (inset) 5 µm. (**F**) 3D colocalization analyses were performed using the JACoP Fiji plugin, and results are displayed as Tukey boxplots (three biological repeats, n> 38 image stacks per condition). The whiskers (shown here as vertical lines) extend to data points no further from the box than 1.5× the interquartile range. For all panels, the P-values were determined using a randomization test. NS indicates no statistical difference between the mean values of the highlighted condition and the control.

**Fig. S5.**
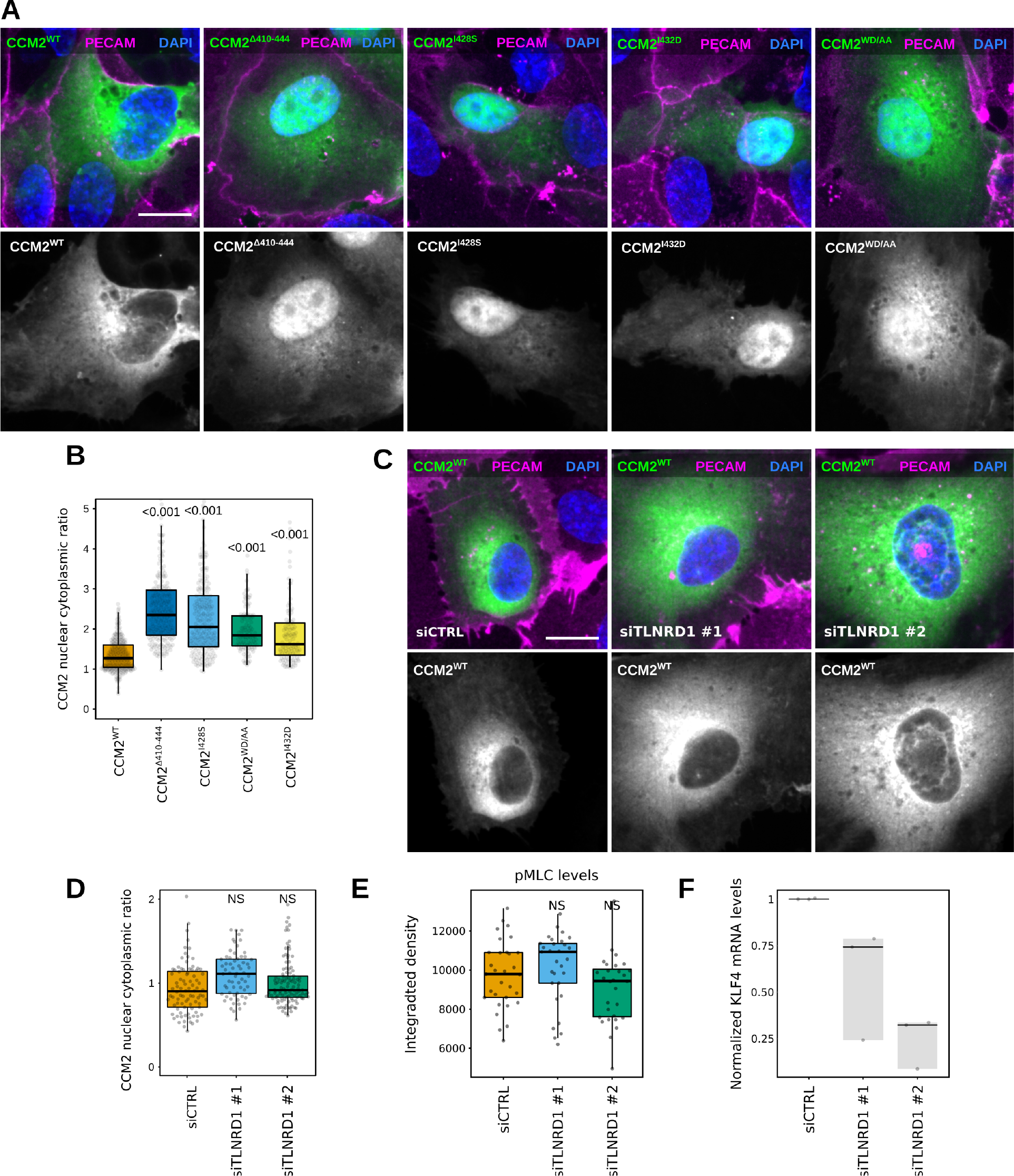
TLNRD1 does not regulate CCM2 localization or function. (**A**-**B**) HUVECs expressing various CCM2-GFP constructs were stained for DAPI and PECAM and imaged using a spinning disk confocal microscope. (**A**) SUM projections are displayed. Scale bar: 10 µm. (**B**) For each condition, the CCM2 nuclear-cytoplasmic ratio was quantified (three biological repeats, n> 110 cells per condition). (**C**-**F**) TLNRD1 expression was silenced in HUVECs using two independent siRNA. (**C**-**D**) Cells were then transfected to express CCM2-GFP and allowed to form a monolayer. Cells were fixed and stained for DAPI and PECAM and imaged using a spinning disk confocal microscope. (**C**) SUM projections are displayed. Scale bar: 10 µm. (**D**) For each condition, the CCM2 nuclear-cytoplasmic ratio was quantified (three biological repeats, n> 60 cells per condition). (**E**) HUVECs were allowed to form a monolayer without flow stimulation. Cells were then fixed and stained for phospho-Myosin light chain (pMLC S20) before being imaged on a spinning disk confocal microscope. The overall integrated density was measured for each field of view from SUM projections (three biological repeats, n = 45 FOV per condition). (**F**) HUVECs were then allowed to form a monolayer without flow stimulation. KLF4 expression levels were measured by qPCR. (**B, D**, and **E**) The results are displayed as Tukey boxplots. The whiskers (shown here as vertical lines) extend to data points no further from the box than 1.5× the interquartile range. For all panels, the P-values were determined using a randomization test. NS indicates no statistical difference between the mean values of the highlighted condition and the control.

**Fig. S6.**
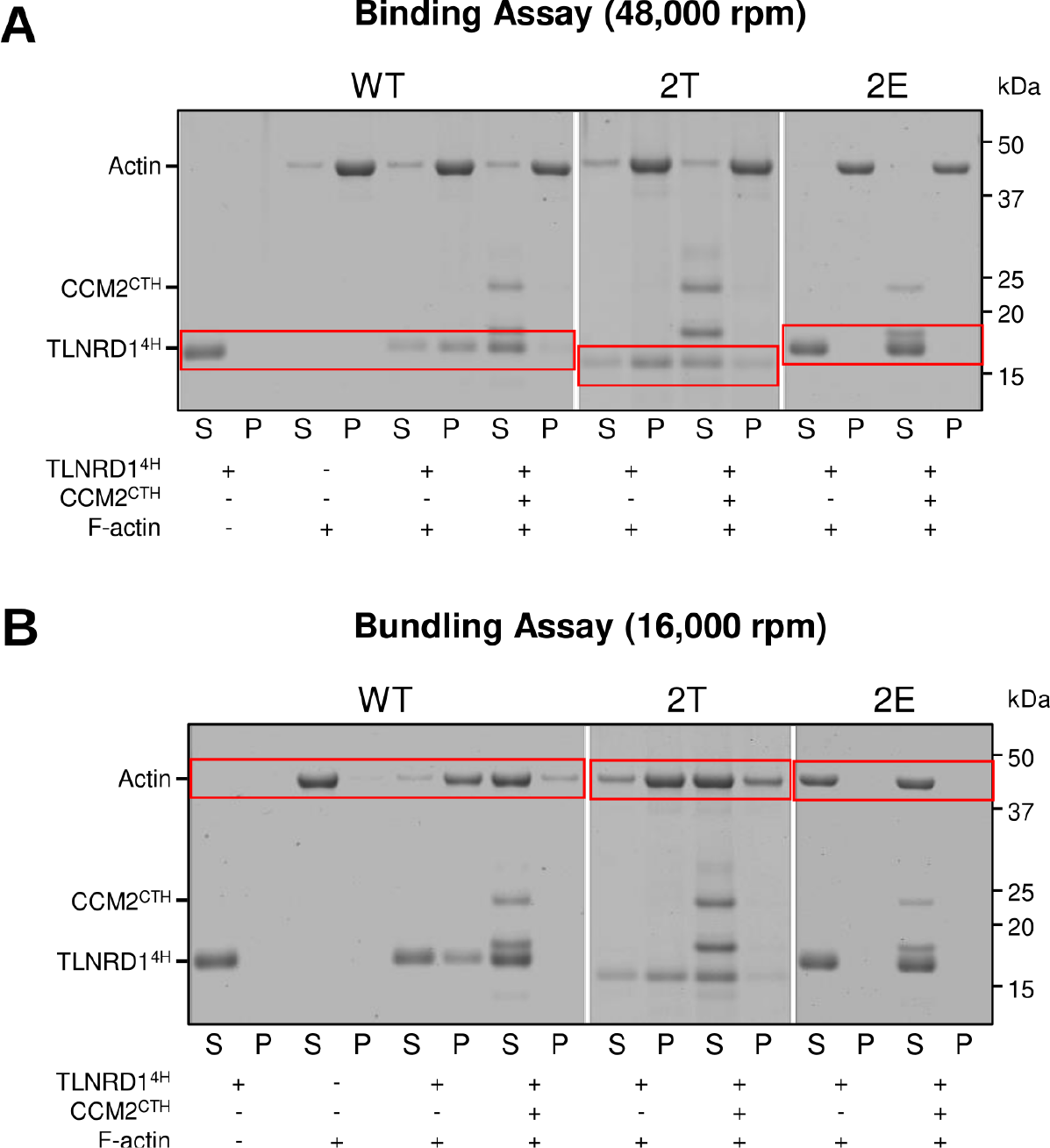
CCM2 inhibits TLNRD1 - actin binding and bundling. (**A**-**B**): Actin co-sedimentation assay with TLNRD1^4H^ mutants in the presence or absence of CCM2^CTH^. Centrifugation at high (**A**, 48,000 rpm) and low (**B**, 16,000 rpm) speeds can distinguish between the F-actin bundling and binding capability of the tested protein constructs. In the absence of CCM2^CTH^, TLNRD1^4H^ both binds and bundles F-actin. The addition of CCM2^CTH^ prevents TLNRD1^4H^ from bundling and binding F-actin. The TLNRD1^2T^ mutant partially restores the binding and bundling activity of TLNRD1 as it interacts only weakly with CCM2^CTH^. The TLNRD1^2E^ mutant cannot bind or bundle F-actin, even without CCM2^CTH^.

